# Glacial history and landscape features shape the hierarchical population genetic structure of woodland caribou (*Rangifer tarandus caribou*) in western Canada

**DOI:** 10.1101/2025.10.15.682449

**Authors:** Samuel Deakin, Anita Michalak, Maria Cavedon, Charlotte Bourbon, Margaret M. Hughes, Lalenia Neufeld, Agnès Pelletier, Jean Polfus, Helen Schwantje, Robin Steenweg, Caeley Thacker, Madeline Trottier, Marco Musiani, Jocelyn Poissant

**Affiliations:** Faculty of Veterinary Medicine, University of Calgary, Calgary, AB T2N 1N4, Canada; Department of Biological Sciences, University of Calgary, Calgary, AB T2N 1N4, Canada; Jasper National Park of Canada, Parks Canada, Jasper, Canada; British Columbia Ministry of Water, Land, and Resource Stewardship, Terrestrial Species Recovery Branch, Prince George, BC, Canada; Canadian Wildlife Service – Pacific Region, Environment and Climate Change Canada, Kelowna, BC V1V 1V9, Canada; Department of Biology, University of British Columbia, Kelowna, BC, Canada; (emeritus) Wildlife and Habitat Branch, Ministry of Water, Lands, and Natural Resource Stewardship, Government of British Columbia, Nanaimo, BC, V9T 6JT, Canada; Wildlife and Habitat Branch, Ministry of Water, Lands, and Natural Resource Stewardship, Government of British Columbia, Nanaimo, BC, V9T 6JT, Canada; Dipartimento Scienze Biologiche Geologiche Ambientali, Università di Bologna, Bologna, Italia

**Keywords:** conservation genomics, population structure, genetic differentiation, glacial history, evolutionarily significant units, endangered species

## Abstract

Caribou, listed as a species at risk across Canada, have experienced a wide range of evolutionary and selective pressures at multiple scales, from large-scale range shifts and recolonisations driven by glacial cycles to more localized contemporary habitat degradation and fragmentation. Given these multi-scale evolutionary forces, genetic variation and diversity are expected to be hierarchically structured. Characterising hierarchical population structure is crucial to understanding a species’ evolutionary history and informing effective conservation and management strategies. In this study, we analysed genomic diversity and variation in woodland caribou (*Rangifer tarandus caribou*) across western Canada using genotypes from ∼33,000 Single Nucleotide Polymorphism (SNP) loci from 759 geo-referenced individuals spanning 45 pre-defined subpopulations. We employed genetic clustering methods and measures of genetic differentiation to characterize hierarchical population structure in the region and tested for latitudinal changes in heterozygosity resulting from post-glacial recolonisation and hybridisation. Our results confirm that woodland caribou genetic diversity and differentiation occur at multiple hierarchical levels, reflecting post-glacial recolonisation patterns and landscape heterogeneity. Notably, the major genetic clusters identified in our study do not align with current conservation units for the species in this region. We also observe elevated heterozygosity in the mid-latitudes of the sampled range, indicative of hybridisation following secondary contact during post-glacial recolonisation. These findings underscore the need to consider and include genetic diversity at all hierarchical levels in conservation planning, as wide-ranging species often experience diverse and complex evolutionary histories and pressures.

## Introduction

The population genetic structure of a species provides valuable insights into both its historical and contemporary dynamics (Hewitt, 2004; Shafer et al., 2010). Patterns of genetic differentiation such as variation in diversity and connectivity across a species’ range, can reveal past colonisation and recolonisation events (Deakin et al., 2020; Shafer et al., 2011; Stone & Cook, 2000). Additionally, these patterns inform our understanding of contemporary gene flow, shedding light on the biotic and abiotic factors that shape genetic connectivity (Breistein et al., 2022; Epps et al., 2018; Forbes & Hogg, 1999). Understanding the natural history and contemporary gene flow of a species can be crucial for its management and conservation, particularly in defining and prioritising conservation units below the species level (Crandall et al., 2000; Fraser & Bernatchez, 2001; Hoelzel, 2023; Moritz, 1994).

Often, the processes that cause population structure do not act in isolation and can occur at different temporal and spatial scales, leading to a phenomenon known as hierarchical population structure (Vähä et al. 2007). For instance, some anadromous fish are structured by both major rivers and sub-structure within water catchments (Poissant et al., 2005; Vähä et al., 2007). Similarly, terrestrial species often show major discontinuities in population structure, due to past glacial refugia (Shafer et al. 2010), and finer substructure due to localized geographic features such as rivers, mountains, and differing habitat types (Cross et al., 2016a; Deakin et al., 2020; Jenkins et al., 2018; Sim et al., 2019). Identifying nested population subdivisions can be challenging for species with broad distributions due to the complex interactions of historical and ecological processes, as well as the need for conducting adequate sampling across vast geographic areas (Cheeseman et al., 2019; Warnock et al., 2010). However, recognizing hierarchical structure is essential for defining conservation units, as it helps identify and delineate evolutionary significant lineages which may be obscured when only considering local or broad-scale differentiation (Cross et al., 2016b; Warnock et al., 2010).

In the conservation of species, the characterisation of unique genetic, ecological, or behavioural adaptation below the species level can help identify populations upon which to focus conservation efforts (Crandall et al., 2000; Fraser & Bernatchez, 2001; Hoelzel, 2023; Moritz, 1994; Ryder, 1986). In the United States conservation units are often identified, managed, and protected as Evolutionarily Significant Units, which from a genetic standpoint are used to identify, delineate, and preserve groups of animals or plants containing unique and putatively-adaptive genetic variation (Crandall et al., 2000; Fraser & Bernatchez, 2001; Hoelzel, 2023; Moritz, 1994). Similarly, in Canada, conservation units are identified and managed as Designatable Units (DUs) listed under the Species at Risk Act (SARA). These DUs are intended to capture unique, significant, and irreplaceable components of Canada’s biodiversity (COSEWIC, 2011; Harding, 2022; Muir et al., 2021).

Caribou (*Rangifer tarandus*) and their conspecifics, reindeer, have a circumpolar distribution (Flagstad & Røed, 2003; Yannic et al., 2014) and are divided into subspecies that inhabit diverse regions and habitat types (Harding, 2022). Like many other terrestrial species, caribou recolonised northwestern North America following the last glacial maximum from at least two refugia; the Beringian refugium, located in northeastern Siberia and northwestern North America, and the North American refugium, consisting of much of the present-day mainland United States (Shafer et al., 2010; Hewitt, 2004; Flagstad & Røed, 2003; Yannic et al., 2014). Woodland caribou (*R. t. caribou,* Banfield 1961) inhabit the southern regions of the species’ distribution in North America and stem from two lineages reflective of their glacial history; the Beringian-Eurasian Lineage (BEL) and the North American Lineage (NAL) (Cavedon et al., 2022a; McDevitt et al., 2009; Polfus et al., 2017a; Yannic et al., 2014). Within this subspecies, various ecotypes and subpopulations are observed (Theoret et al., 2022) which exhibit behaviours and adaptations linked to their ancestral lineages (Cavedon et al., 2022b; McDevitt et al., 2009). Given their broad distribution, presence of differing glacial ancestries, and previously identified population structure (McLoughlin et al., 2004; Priadka et al., 2019; Serrouya et al., 2012; Wilson et al., 2022), woodland caribou in western Canada likely exhibit a hierarchical population structure (Michalak 2023).

In Canada, caribou subpopulations, often referred to as “herds”, are grouped into DUs or SARA-listed units for conservation purposes (Weckworth et al., 2018). In western Canada, the focal region of this study, caribou DUs and SARA delineations are in part attributed to both phylogeographic history and population genetic structure (Cronin et al., 2005; Klütsch et al., 2016; McDevitt et al., 2009; Polfus et al., 2017b; Serrouya et al., 2012; Taylor et al., 2021; Weckworth et al., 2012; Yannic et al., 2014); however, many previous studies have used different sets of genetic markers and examined different subpopulations, making it difficult to compare results. At present, four DUs and three SARA-listed units (which overlap the DUs) have been identified in this region (COSEWIC, 2011; SARA, 2012a, 2012b, 2014). In both classification schemes, boreal, northern mountain, and southern mountain populations are differentiated, with additional partitioning within the southern mountain population (Figure 1).

**Figure 1.**
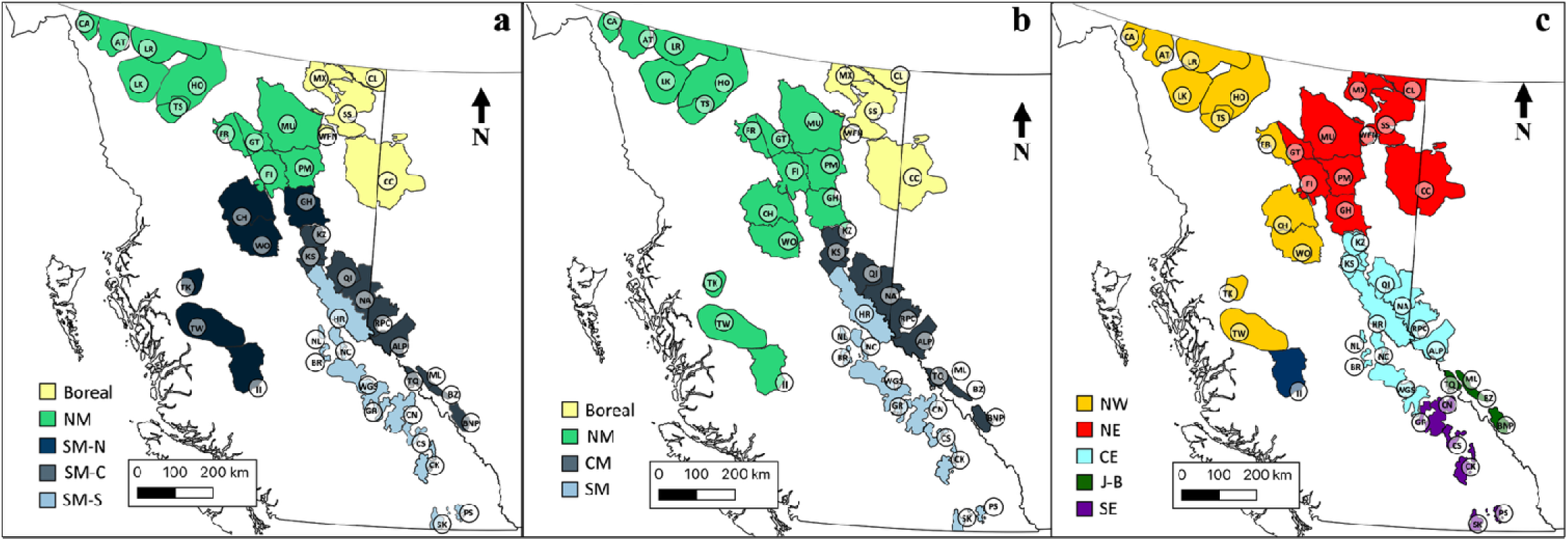
Distribution of studied woodland caribou subpopulations in western Canada, with labels of genotyped subpopulations following the abbreviation scheme in Table 1. (**a**) Depicts classification of subpopulations according to the Species at Risk Act (SARA) Schedule 1 populations: Boreal population, Northern Mountain (NM) population, and Southern Mountain population (SM) Northern (-N), Central (-C) and Southern (-S) groups. (SARA, 2014) **(b)** Depicts Designatable Units (DUs) (COSEWIC, 2011) classification proposed in 2014 by the Committee on the Status of Endangered Wildlife in Canada: Boreal population (DU6), Northern Mountain population (NM; DU7), Central Mountain population (CM; DU8), and Southern Mountain population (SM; DU9).(c) Depicts major genetic clusters inferred from our analyses. Note: here the Frog population is classified as a northwestern population as it is situated west of the Rocky Mountain Trench despite its minor-majority assignment to the northeastern cluster (an overall assignment rate of 50.3%)Given that this subpopulation contained near equal portions of northeastern and northwestern genetic backgrounds, the Frog subpopulation likely represents a transition zone between the two northern clusters. Maps created in QGIS.

**Table 1.**
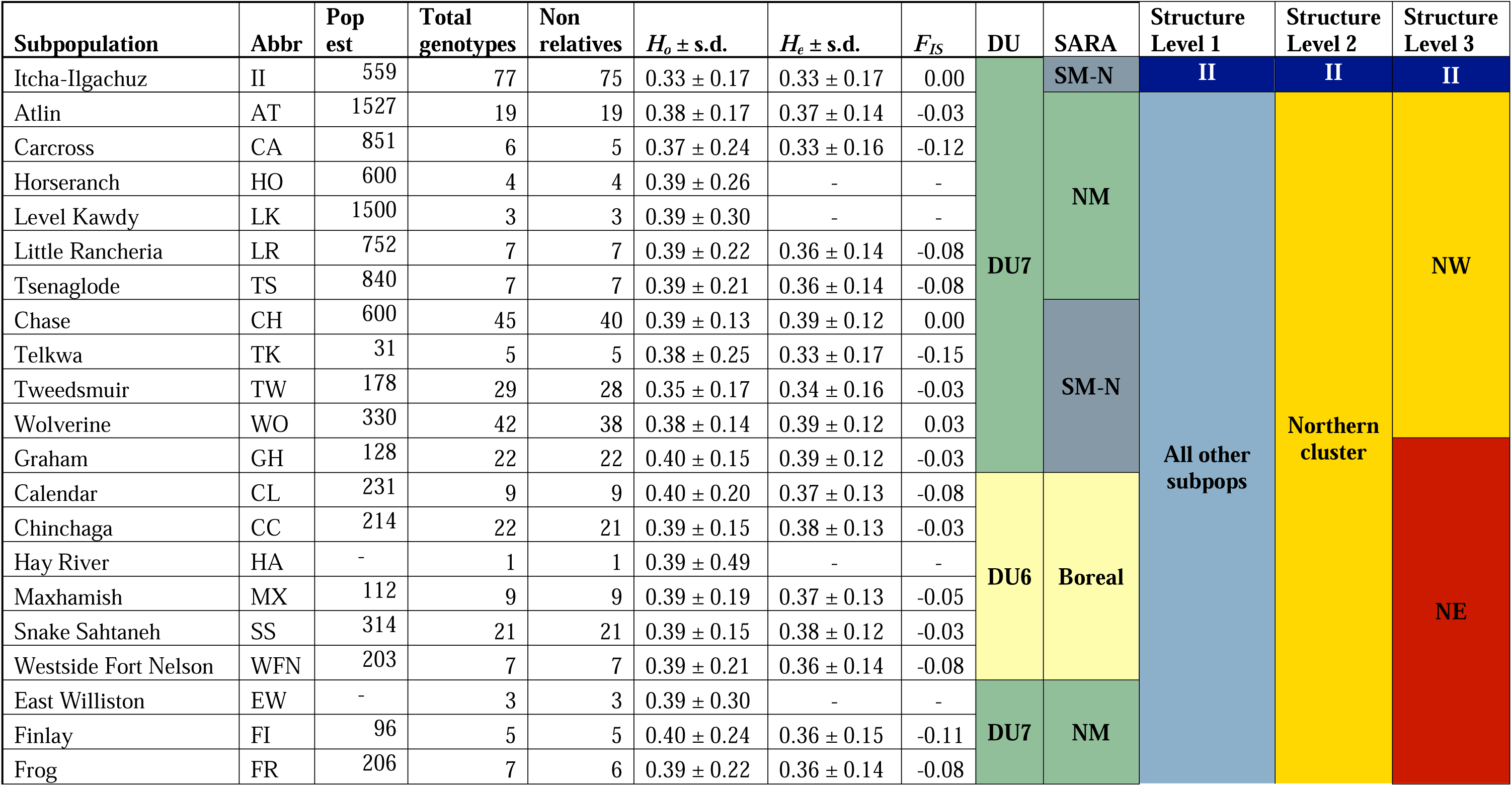
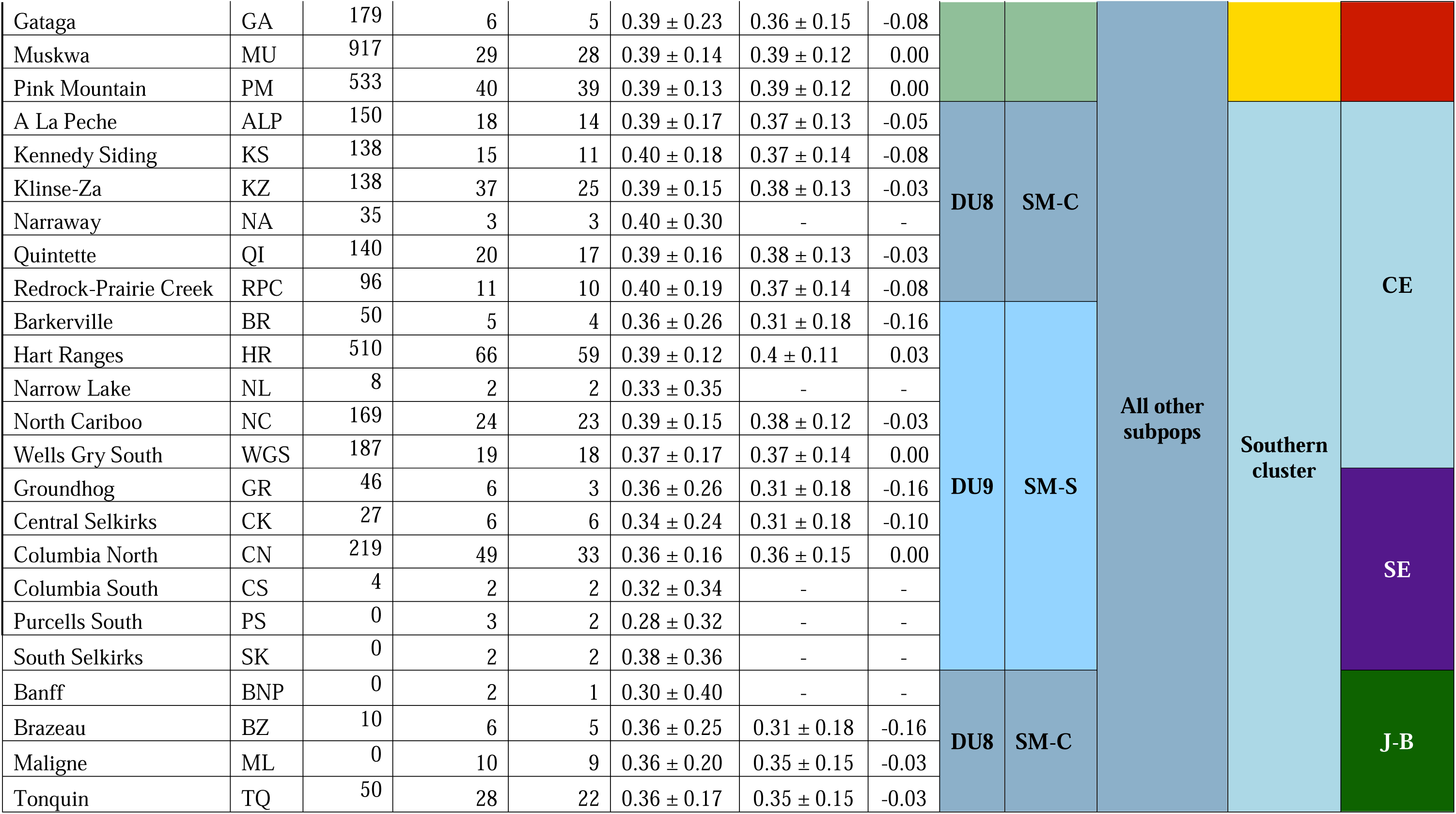
Genetic variation across woodland caribou subpopulations in western Canada. Total number of genotypes remaining after quality filtering (*n*=759), and excluding close relatives (Non-relatives, *n*=678) are provided for each subpopulation, along with observed heterozygosity (*H_o_*), expected heterozygosity (*H_e_*), and *F_IS_* calculated while including close relatives. *H_e_* and *F_IS_* were only estimated for subpopulations containing at least 5 quality-filtered samples. Each subpopulation’s, Designatable Unit (DU), and Species at Risk Act (SARA) classifications, as well as the major genetic cluster it assigned to in hierarchical *Structure* analyses (down to level 3, as depicted on Figures 4 and 5) are also presented. SM-N, SM-C, and SM-S are abbreviations for Southern Mountain-northern group, Southern Mountain-central group, and Southern Mountain-southern group, respectively. II, NW, NE, CE, J-B, and SE are abbreviations for Itcha-Ilgachuz, northwestern, northeastern, central-eastern, Jasper-Banff, and southeastern, respectively. Population estimates (Pop est) taken from last available estimates (COSEWIC, 2014; Government of Alberta, 2017; Government of British Columbia, 2023; Parks Canada, 2018, 2024).

In this study, we build on the preliminary work of Michalak (2023) to conduct a comprehensive hierarchical analysis of genetic diversity and variation in woodland caribou across western Canada. Specific aims were to: (i) identify broad- and fine-scale population genetic structure and its causes, and (ii) test if genetic diversity was higher in the central region of our sampling range due to secondary contact and hybridisation between glacial lineages. To do this we used genotypes from ∼33,000 loci generated using a caribou-specific Single Nucleotide Polymorphism (SNP) array. We expected to find hierarchical population genetic structure concurrent with evolutionary processes at different levels. Specifically, we expected to find two overarching clusters representing differing glacial ancestries, below which we expected landscape features to explain further sub-structure. As for genetic diversity, we expected it would be highest in the middle of the studied range, where the two glacial linages of caribou are known to have hybridised. Given the preliminary results of Michalak (2023), we anticipated genetic structure would not fully align with current management schemes. This study provides insights into the natural history, contemporary gene flow, and factors affecting the population genetic structure of caribou in western Canada, offering valuable guidance for defining conservation units below the species level in this species and other species with similar evolutionary histories.

## Materials and methods

### Samples and geographic data

Woodland caribou blood and tissue samples were collected by Provincial Government and Parks Canada partners in British Columbia and Alberta, Canada, between 2012–2023. Due to the sampling regimes of these partners, most samples came from female animals. All samples were associated with a georeferenced sampling location and/or range of origin.

### DNA extraction and genotyping

DNA was prepared following the protocols described in Michalak (2023). In brief, DNA was extracted using a QIAGEN DNeasy Blood & Tissue or QIAamp 96 DNA QIAcube HT Kit with recommended manufacturer protocols and eluted in 400µL of molecular grade water. DNA was then quantified using either a BioTek Synergy LX Multimode Reader or Thermo Fisher Qubit 4 Fluorometer and the Thermo Fisher Quant-iT and Qubit dsDNA Assay Kits, respectively. Eight hundred and fifty-four samples from 45 pre-defined subpopulations (Table 1, Figure 1) containing ≥ 400 ng of DNA were normalized to a quantity of 400 ng, dried on a Thermo Scientific Savant SpeedVac DNA 130 Integrated Vacuum Concentrator System, and sent to the Centre d’expertise et de service Génome Québec (Montreal, Canada) for genotyping using an Illumina single nucleotide polymorphisms (SNP) array targeting ∼60,000 loci distributed across the caribou genome (Carrier et al., 2022).

### Loci mapping and genotype filtering

Positional data for the SNPs were originally provided for the ULRtarCaribou_2 genome scaffold-level assembly (GCA_019903745.1) (Prunier et al., 2022). However, for our analysis we opted to remap these SNPs to the second version of this genome, ULRtarCaribou_2v2 genome (GCA_019903745.2), a chromosomal-level assembly (Poisson et al., 2023). To do so, we aligned the source sequence data from the Illumina array manifest file (Supplementary Material 1) to the ULRtarCaribou_2v2 genome assembly (Poisson et al., 2023). First we removed 13,445 uninformative or suboptimal loci as recommended by Carrier et al. (2022) (see Supplementary Materials of Carrier et al. (2022) for the list of loci). Each SNP’s source sequence was then formatted as an entry in a FASTA file and aligned to ULRtarCaribou_2v2 using bowtie2 v2.3.1 (Langmead & Salzberg, 2012) and the *sensitive* alignment parameter. Samtools v1.6 (Li et al., 2009) was then used to call SNPs and obtain positions (see Supplementary Information 1 for code and parameters).

Genotype filtering was performed as described in Michalak (2023). In brief, SNP genotypes received from the Centre d’Expertise et de Service Génome Québec were filtered using PLINK v1.9. First, we filtered for mapped SNPs (described above). Then a few samples identified as duplicates were removed using a > 95% similarity threshold and information from the PLINK *genome* function. The PLINK functions *list-duplicate-vars* and *exclude* were then used to identify and remove duplicate SNPs. Further filtering consisted of excluding individuals with < 95% genotyping rate (*mind* 0.05), as well as SNPs with < 95% genotyping rate (*geno* 0.05), deviating from Hardy-Weinberg equilibrium (*hwe* 1e-6), with a minor allele frequency < 0.05 (*maf* 0.05), and in strong linkage disequilibrium with another SNP (*indep-pairwise* 50 5 0.5). Ultimately, 759 individuals from the 45 pre-defined subpopulations (Table 1) and 32,440 SNPs were retained. A dataset excluding 81 close relatives (parent-child or full sibling relationships, where one individual from each pair was retained) was also generated for use in specific analyses. This was done with the same filtering pipeline following the exclusion of close relatives using the PLINK v2 (Chang et al., 2015) *king-cutoff* command with a threshold of 0.177. This dataset contained a total of 678 individuals and 32,636 SNPs.

### Genetic diversity

To examine patterns of genetic diversity across the study area, we used the R (R Core Team, 2013) package *dartR* v2.9.7 (Mijangos et al., 2022) to calculate observed heterozygosity (*H_o_*) of each pre-defined subpopulation. We also calculated expected heterozygosity (*H_e_*) for each pre-defined subpopulation with at least 5 genotyped individuals and inferred genetic cluster. These were then used to calculate *F_IS_* based on the formula (*H*_e_ – *H*_o_)/*H*_e_ (Wright, 1949). We also estimated the degree of differentiation between pairs of pre-defined subpopulations as well as inferred genetic clusters for which both expected and observed heterozygosity was available. This was accomplished using pairwise fixation index (*F_ST_*) values calculated using *StAMMP* v1.6.3 (Pembleton et al., 2013) in R, with significance assessed using 1,000 bootstraps.

To assess if latitude (a proxy for position between the northern and southern glacial refugia) was associated with heterozygosity, we modelled *H_e_*as a function of latitude and/or recent census size to account for the latter’s likely relationship with heterozygosity. Population estimates were obtained from government reports (COSEWIC, 2014; Government of Alberta, 2017; Government of British Columbia, 2023; Parks Canada, 2018, 2024) and are reported in Table 1. The last population estimate for the Maligne subpopulation was 0 (as the subpopulation was extirpated); in this case a population estimate of 5 was used to allow for its inclusion in this analysis. Specifically, we examined a series of generalized linear models (GLM) with a Gaussian error distribution. In these models, we tested all combinations of census size and latitude fitted as different terms as fixed effects (Table S1). Census size was tested as both a linear and log transformed term given that *H_e_* should eventually plateau as census size increases. Latitude was tested as both a linear and quadratic term given that we expected higher subpopulation *H_e_* values in the middle of our range where hybridisation has taken place. To select the best fitting model, we examined the *r^2^* values of the models and the Akaike Information Criterion corrected for small sample size (AIC_C_). These analyses were conducted in R using the *glm* function from the *stats* v4.2.1 package. Data visualization was performed using *ggplot2* v3.5.1 (Wickham, 2011) and *Visreg* v2.7.0 (Breheny & Burchett, 2017).

### Population genetic structure

We assessed population genetic structure using a combination of model- and distance-based approaches. For these analyses we excluded close relatives. As an initial assessment, we conducted a Principal Component Analysis (PCA) and a Discriminant Analysis of Principal Components (DAPC) using *adegenet* v2.1.10 (Jombart, 2008; Jombart & Ahmed, 2011) in R. The *find.clusters* function was used to examine all principal components (PCs) and identify the best-fitting number of clusters given Bayesian Information Criterion (BIC) values for *K* clusters ranging from 1 to 45. We interpreted the best number of clusters as being the point where the curve of BIC values as a function of *K* elbowed (Jombart & Collins 2015; Thia 2023). To describe the clusters identified using DAPC we chose to retain all eigenvalues for *K*-1 discriminant functions, as well as 44 PCs to match the number of pre-defined subpopulations minus one, as recommended by Thia (2023). Finally, we incorporated the first two discriminant functions in a scatterplot to visualize variation among identified groups.

Population structure was further evaluated using the Bayesian clustering approach implemented in *Structure* v2.3.4 (Pritchard et al., 2000), which groups individual genotypes into *K* clusters that maximize within-cluster Hardy-Weinberg and linkage equilibria. Because genetic structure likely occurs at multiple levels in woodland caribou we performed *Structure* analyses in a hierarchical fashion (see Vähä et al., (2007) for a description of this approach). We initially ran *Structure* ten times for each value of *K* from 1– 10 using the admixture model, correlated allele frequencies, and no *a priori* grouping of individuals. Each run consisted of a burn-in of 20,000 iterations followed by 50,000 Markov chain Monte Carlo (MCMC) repetitions, which was assessed as adequate based on convergence (Wang 2022). The *R* package *pophelper* v2.3.1 (Francis, 2017) was then used to calculate the Δ*K* statistic of (Evanno et al., 2005) to examine which value of *K* was best supported by the data. Additionally, *pophelper* was used to consolidate clusters from multiple iterations of *Structure* and visualize results. As in Vähä *et al*. (2007), we then ran additional *Structure* analyses for each of the clusters identified using the same parameters as above. As the Δ*K* statistic cannot determine the presence of only one true genetic cluster (Janes et al., 2017), we visually examined Q-matrices assignments to determine at which hierarchical level analyses should stop. Given our research objective of characterising broad-scale genetic structure, we ceased analyses when clusters were being identified within pre-defined subpopulations, as clusters below the subpopulation level likely represent family groups.

Population structure was also examined using the spatially explicit R program *TESS3* v1.0 (Caye et al., 2016) implemented in R. In contrast to *Structure*, *TESS* assigns individuals to clusters while incorporating information on each sample’s geographic location. In instances where samples were not associated with a precise sampling location, we assigned a sampling location in one of two ways. If precise sampling locations for other individuals from the same pre-defined subpopulation was available, we assigned a random GPS location around the centroid of these known sampling locations within the maximum extent of the distribution of other subpopulation members. When precise locations were not known for any individual from the subpopulation, we generated random locations around a putative subpopulation central location, with the maximum extent of the distribution set as the average outer limit observed across all pre-defined subpopulations with precise individual capture locations. For the *TESS3* analysis we conducted 10 runs for values of *K* ranging from 1 to 45 (tolerance = 1×10^-7^, max. iterations = 1,000) and used the cross-entropy criterion to select the optimal value of *K*, which corresponds to the one with the lowest cross-validation score. We also used the *tess3r* v1.1.0 (Caye et al., 2016) R package to create maps of the geographic distribution of genetic clusters and their corresponding geographic boundaries.

We constructed an individual-based neighbor-joining tree using Manhattan distances from a genotype matrix calculated using the R package *BEDMatrix* v2.0.4 (Grueneberg & de los Campos, 2019). The neighbor-joining tree was inferred using the *nj* function in the *ape* v5.8 (Paradis et al., 2004; Paradis & Schliep, 2019) R package and its confidence assessed using 1,000 bootstraps. The tree was then visualized using the *ggtree* v3.15.0 R package (Yu et al., 2017).

### Isolation-by-distance

To examine how spatial separation influenced trends in genetic differentiation between subpopulations we tested for patterns of isolation-by-distance across the study area. We examined if Euclidean geographic distance between the centroid of capture locations for each subpopulation with at least five genotyped individuals was associated with Nei’s genetic distance (Nei, 1972, 1978) using a Mantel test with the R package *ade4* v1.7.22 (Chessel et al., 2004; Thioulouse et al., 1997). Since population structure can skew isolation-by-distance results when using Mantel tests (Meirmans, 2012), we also performed separate tests for each major genetic cluster inferred at levels 2 and 3 of our hierarchical analysis.

### Isolation by landscape features

To investigate the effect of specific landscape features on genetic differentiation, we used partial Mantel tests to assess whether features identified as boundaries in our clustering analysis influenced genetic distances between subpopulations, while controlling for geographic distance. We calculated Nei’s genetic distances between subpopulations, geographic distances, and constructed a binary matrix indicating whether subpopulation pairs were on the same (0) or opposite sides (1) of a given landscape feature. Only subpopulations with greater than five genotyped individuals were included in this analysis. All partial Mantel tests were conducted using the *vegan* package v2.7.1 in R.

## Results

### Genetic diversity within and among populations

Overall *H_o_* and *H_e_* were 0.38 ± 0.10 (s.d.) and 0.41 ± 0.10 and ranged from 0.28–0.40 and 0.31–0.40 within pre-defined subpopulations, respectively (Table 1). Mean pairwise *F_ST_*between all subpopulations was 0.08 (range -0.0005–0.18), with the greatest values observed between the Itcha-Ilgachuz and Brazeau subpopulations, and the lowest between Finlay and Pink Mountain subpopulations (Supplementary Material 2). *F_IS_* ranged from -0.16 to 0.03 (Table 1). Notably, most *F_IS_* values were negative, likely a result of gene flow between subpopulations given the low pairwise *F_ST_* values observed across the sampling range. Genetic diversity varied across the sampling range, being highest in the center and decreasing toward higher and lower latitudes. From our analysis of how latitude and population size affect *H_e,_*the best fitting model (model 11; *r^2^* = 0.56, AIC_C_ weight = 0.669) was supported over the next best model by a ΔAIC_C_ of 1.86 (Supplementary Table S1). In the best fitting model, *H_e_* increased with the linear term for latitude, decreased with the quadratic term for latitude, and showed a near significant positive relationship with log-transformed census size (Figure 2, Supplementary Figure S1, Table S2).

**Figure 2.**
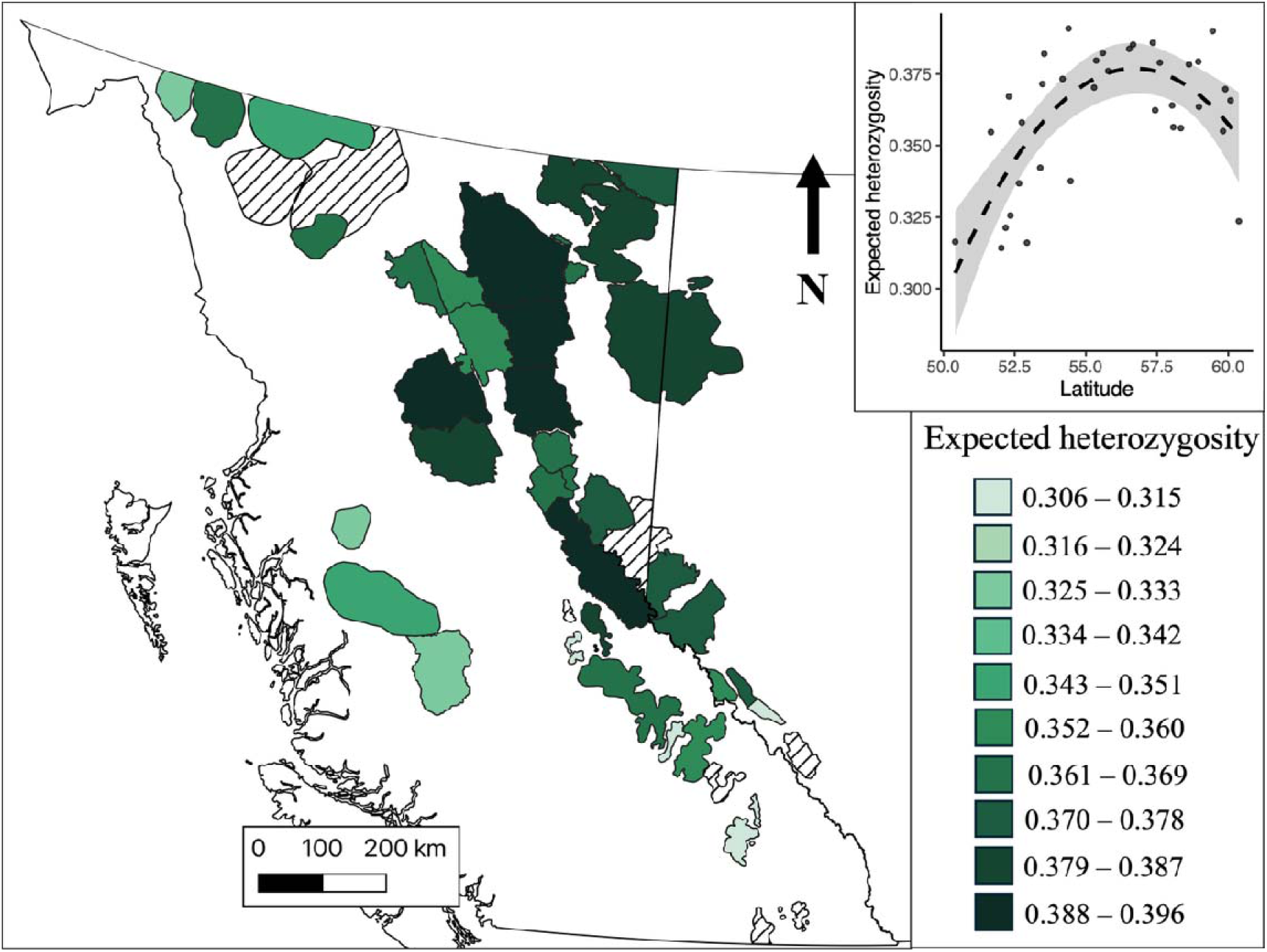
Expected heterozygosity across Woodland caribou subpopulations in western Canada (n=45). Crosshatching represents subpopulations with less than five individuals genotyped, and outlines with no colour represent unsampled subpopulations. Insert in top right was generated using the R *visreg* function and shows the predicted association between latitude and expected heterozygosity from the best fitting linear model. The grey area depicts 95% confidence intervals.

### Population genetic structure

A PCA of all individuals suggested the presence of multiple genetic clusters across the study range (Figure 3). The first PC, which explained 3.45% of the variation, mostly distinguished individuals belonging to the Itcha-Ilgachuz and Tweedsmuir subpopulations of western British Columbia (BC) from all other subpopulations. The second PC, which explained 2.64% of the variation, primarily separated individuals found in the northern part of the sampled range from those found in the central and southern parts. The DAPC indicated the most optimal number of clusters to be between four and seven, and the resulting scatterplots exhibited patterns similar to the PCA (Figure S2).

**Figure 3.**
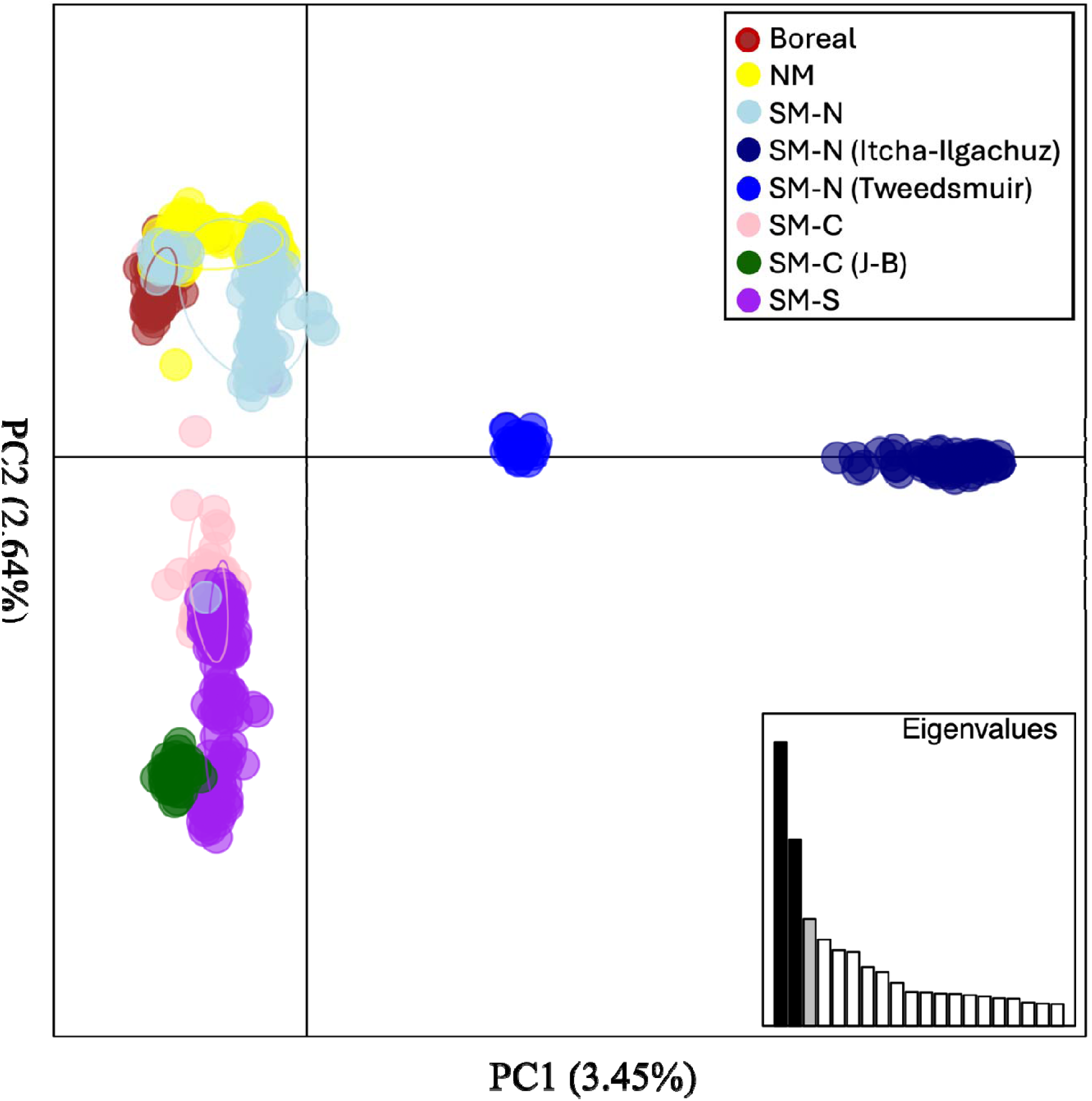
Principal Component Analysis (PCA) of woodland caribou in western Canada. Points in the PCA represent individuals, categorized according to their respective SARA groupings (SARA, 2012b, 2012a, 2014). Itcha-Ilgachuz and Tweedsmuir subpopulations, and Jasper-Banff (J-B) from the SM-N and SM-C groups, respectively, are highlighted to show discontinuities in the groupings. NM represents Northern Mountain.

The hierarchical *Structure* analysis resulted in six levels of clustering. The Δ*K* plot of the initial analysis including all samples indicated *K=* 2 as the most supported number of clusters (Figure S3), with the Itcha-Ilgachuz subpopulation clustering separately from all other subpopulations (Figure 4 [level 1]). The subsequent analysis of all remaining samples (excluding Itcha-Ilgachuz) also indicated *K=* 2 as the most supported number of clusters within this reduced group of samples (Figure S4), which indicated a northern cluster and a southern cluster (Figure 4 [level 2], Table 1). The Δ*K* plots for the resulting northern (Figure S5) and southern (Figure S6) clusters further supported dividing the northern cluster in two; resulting in a northeastern and northwestern, and the southern cluster into three; resulting in a central-eastern, a Jasper-Banff, and a southeastern cluster (Table 1, Figure 4 [level 3], Figure 5).

**Figure 4.**
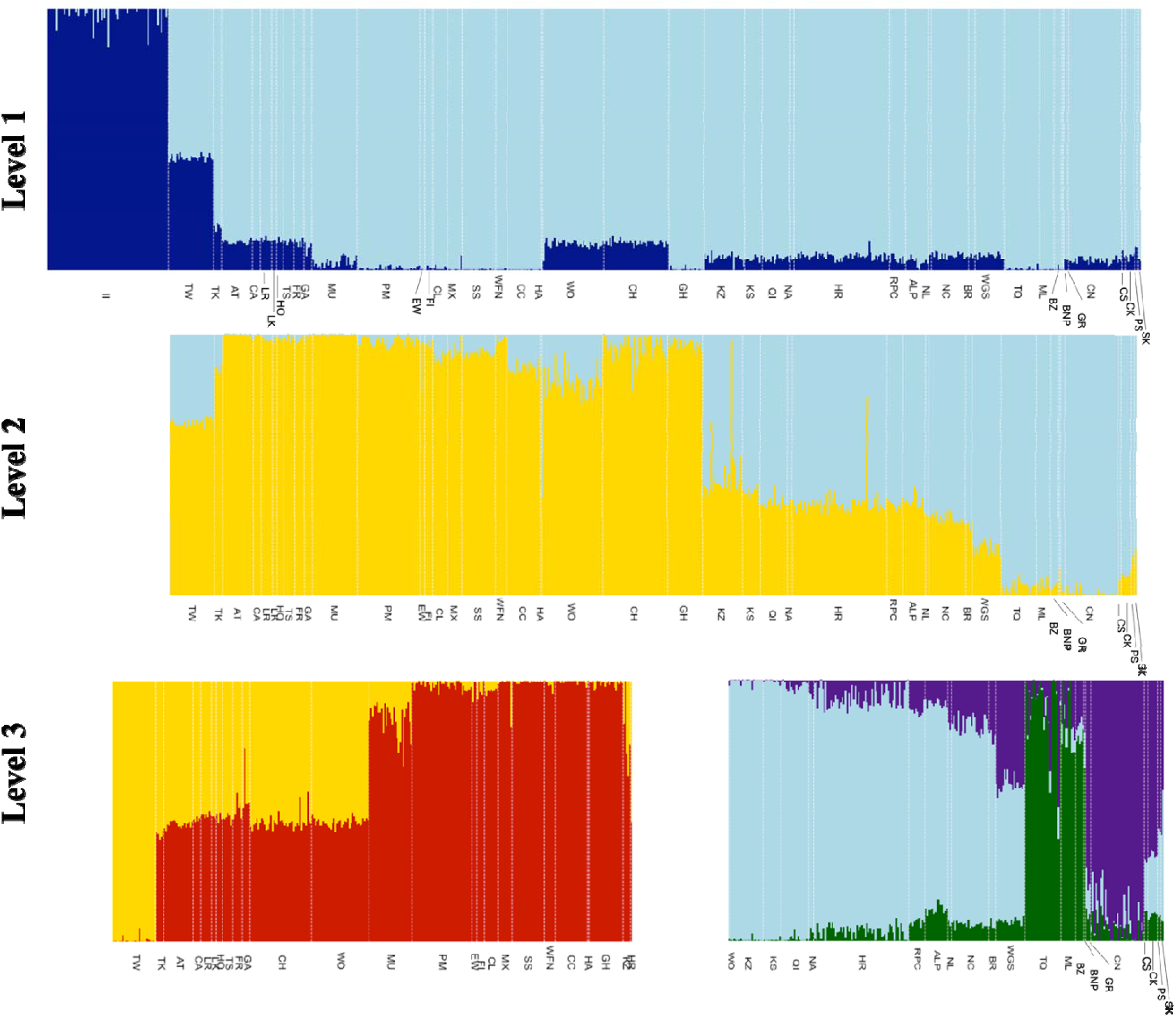
Admixture plots for all woodland caribou included in subpopulation structure analysis (*n=*678), at level 1 all individuals were included in the analysis, at level 2 all individuals except Itcha-Ilgachuz individuals were included in the analysis (dark blue at level 1), at level 3 analysis was run on the two separ te clusters identified at level 2. Labels correspond to subpopulations, for full information on subpopulations see Table 1.

**Figure 5.**
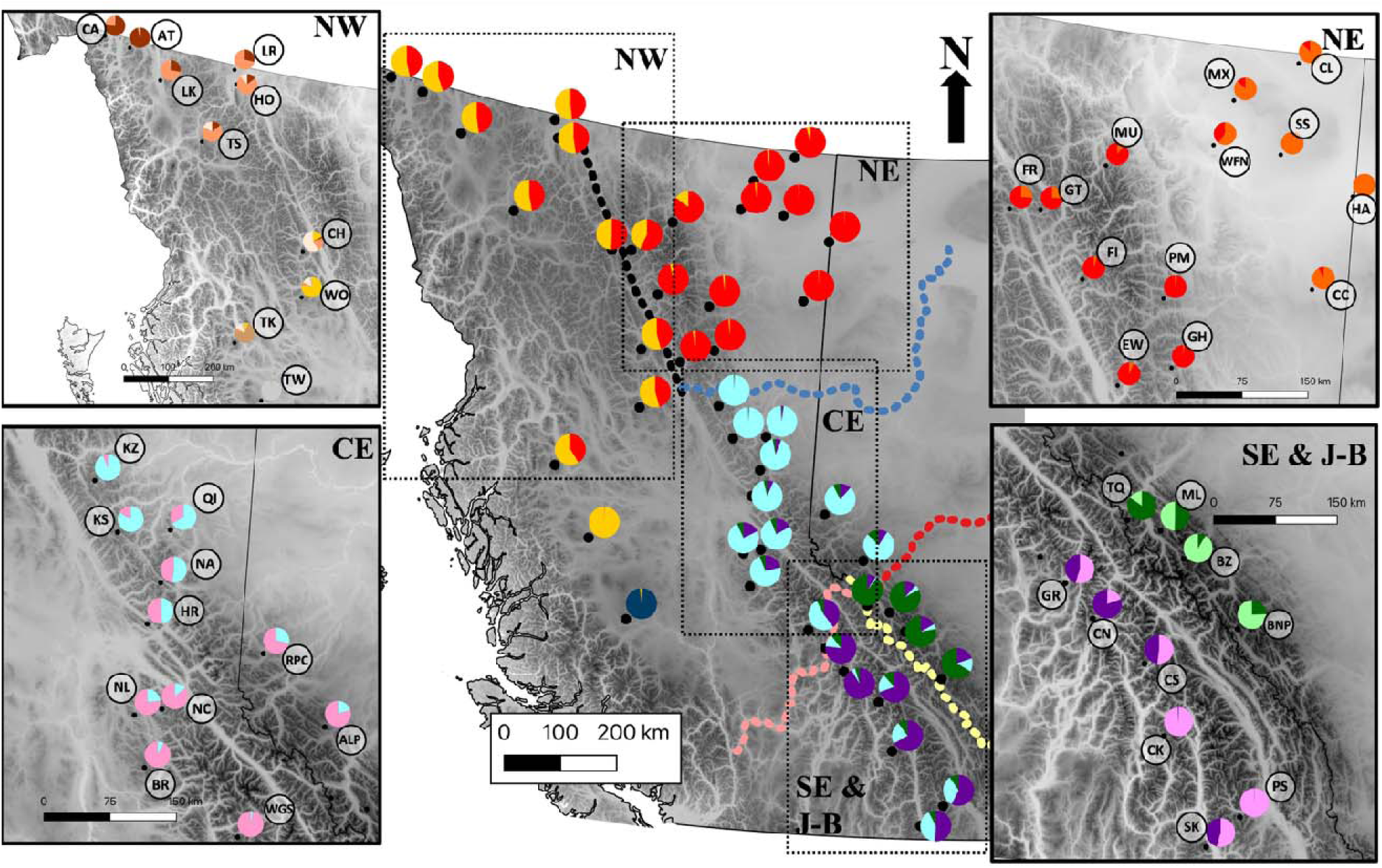
Woodland caribou subpopulations in western Canada and their admixture of each of the genetic clusters identified in *Structure* analysis at different hierarchical levels. Main panel shows the six main genetic clusters identified in western Canada, top left shows genetic clusters identified within the northwestern cluster (NW), top right shows genetic clusters identified within the northeastern cluster (NE), bottom left shows genetic clusters identified within the central-eastern cluster (CE), and bottom right shows genetic clusters identified within the Jasper-Banff (J-B) local population unit cluster and southeastern cluster (SE & J-B). Dashed lines represent landscape features which define cluster boundaries; Rocky Mountain Trench (black), Peace River (blue), Yellowhead Pass and Athabasca River (red), North Thompson Valley (pink), Great Divide (yellow). For subpopulation names, major clusters, and other details see Table 1.

Ultimately, hierarchical clustering analyses beyond the aforementioned six major clusters (Figures 4 and 5) found the 45 predefined subpopulations to separate out into a total of 31 clusters (Supplementary Table 3, Figures S7-S21). These finer-resolution clusters generally consisted of a single pre-defined subpopulation, but in some cases pre-defined subpopulations were grouped together (e.g., Klinse-Za and Kennedy Siding) (Figures S7-S21, Table S3).

In the *TESS* entropy criterion plot, cross-validation values continuously declined up to *K* = 45 (the largest *K* tested; Figure S22). *TESS’*s geographic predictions for values of *K* = 2–8 are presented in Figure S22. These broadly align with the clusters identified by *Structure*, except in the *TESS* analysis the Tweedsmuir subpopulation separates as its own cluster from *K* = 6 prior to separation of the Jasper-Banff cluster at *K=7*.

The neighbor-joining tree closely mirrored population structure results from other analyses, showing a clear distinction between Itcha-Ilgachuz, Southern Mountain-northern group (SM-N), Northern Mountain and Boreal. Southern Mountain-central (SM-C) group and -southern (SM-S) group were separate from all other aforementioned groups but were intermixed with each other. Within these groups, individuals clustered by subpopulation but not by SARA subgroups or COSEWIC DUs. Itcha-Ilgachuz and Tweedsmuir individuals (SM-N) formed a distinct branch, separate from all other Northern Mountain, SM-N, and Boreal subpopulations. The remaining SM-N and Northern Mountain individuals were intermixed and appeared on the same branch as all Boreal individuals (which clustered together) (Figure 6).

**Figure 6.**
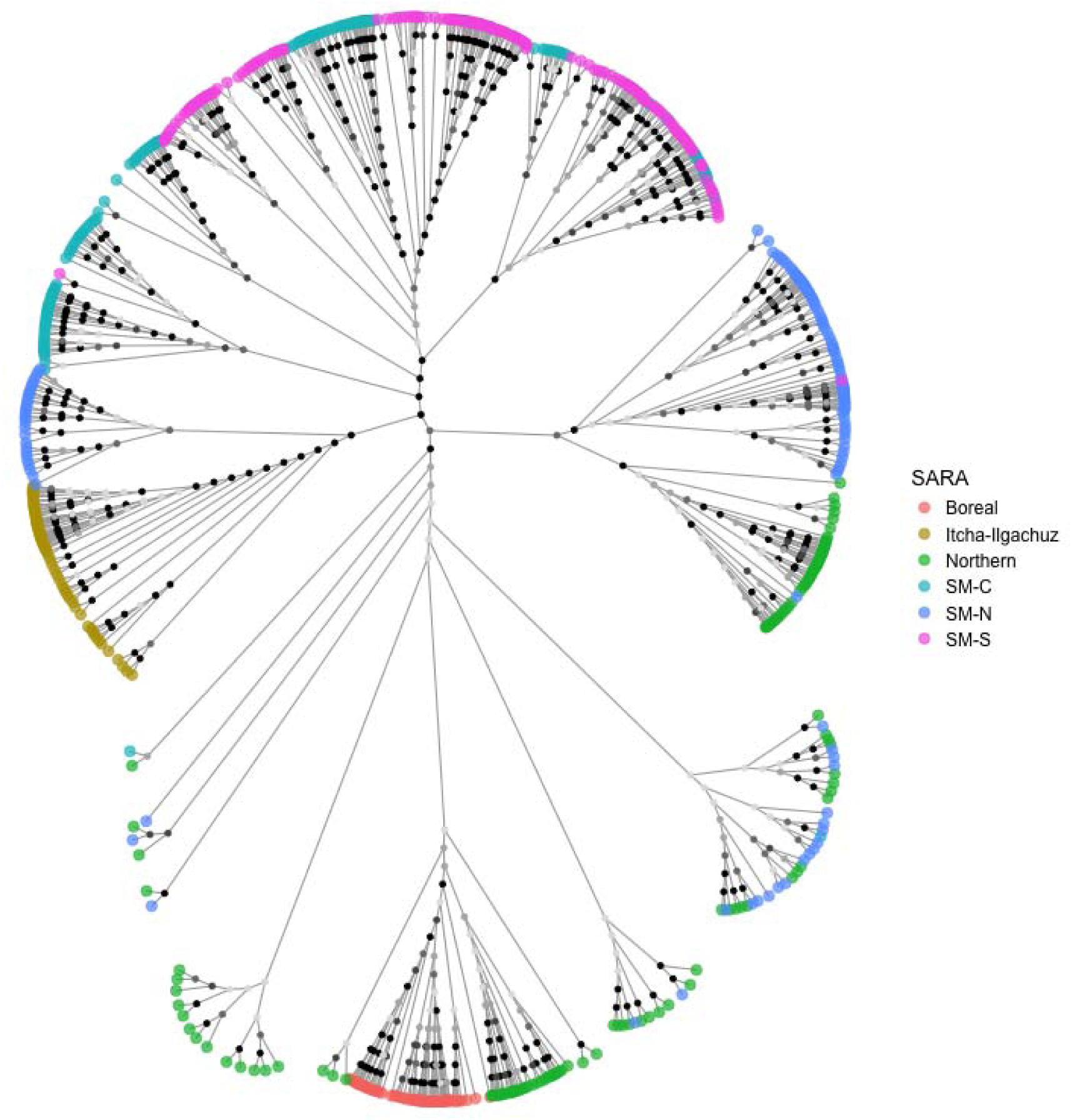
Neighbor-joining tree of woodland caribou sampled throughout western Canada. Branches represent individuals, with tip colours representing each individual’s SARA listing (SARA, 2014) (For a more detailed figure which includes source subpopulation see Figure S27). Bootstrap values were estimated based on 1000 replicates and are represented on internal nodes as circles in five classes/shades of grey (in 20% increments, with the darkest circle representing the 81–100% class). Abbreviations: SM-N = Southern Mountain-northern group, SM-C = Southern Mountain-central group, and SM-S = Southern Mountain-southern group. Note, the SM-N branch beside Itcha-Ilgachuz is composed of all Tweedsmuir individuals.

### Genetic variation within and among major clusters

*H_e_* and *H_o_* within, and *F_ST_*between major clusters were calculated post-hoc after identification during the *Structure* analysis. At the upper hierarchical structure levels (1-3), *H_e_* and *H_o_* ranged from 0.33-0.41 and 0.33-0.38, respectively. Itcha-Ilgachuz had the lowest *H_e_* and *H_o_*, whereas the level 1 main cluster had the highest *H_e_* and the northeastern cluster at level 3 had the highest *H_o_*. Across levels 1-3 *F_ST_* values ranged from 0.02-0.15 and were all significantly different from zero after Bonferroni correction (Table 2). The lowest and highest levels of differentiation were observed at level 3, the lowest between the northeastern and northwestern clusters (*F_ST_* = 0.02), and the highest between the Itcha-Ilgachuz and the Jasper-Banff clusters at level 3 (*F_ST_* = 0.15), with the mean *F_ST_* for level 3 being 0.07. Pairwise *F_ST_* values between clusters at levels 4 to 7 ranged from <0.001 to 0.342 (Supplementary Material 2) while *H_e_* and *H_o_* ranged from 0.19-0.40 and 0.29-0.40 within these clusters (Supplementary Material 3).

**Table 2.** Pairwise *F_ST_* values between inferred genetic clusters for woodland caribou in western Canada. *F_ST_* values and 95% confidence intervals are given below the diagonal, and respective p-values above the diagonal. Cluster labels follow abbreviations defined in Table 1 and presented in Figure 4.

| **Level/cluster** |  |  |  |  |  |  |
| --- | --- | --- | --- | --- | --- | --- |
| **Level 1** | Itcha-Ilgachuz | All other subpopulations |  |  |  |  |
| Itcha-Ilgachuz | – | < 0.001 |  |  |  |  |
| All other subpopulations | 0.083 (0.082-0.084) | – |  |  |  |  |
| **Level 2** | Itcha-Ilgachuz | Northern Cluster | Southern Cluster |  |  |  |
| Itcha-Ilgachuz | – | < 0.001 | < 0.001 |  |  |  |
| Northern Cluster | 0.086 (0.085-0.087) | – | < 0.001 |  |  |  |
| Southern Cluster | 0.095 (0.094-0.096) | 0.024 (0.024-0.024) | – |  |  |  |
| **Level 3** | Itcha-Ilgachuz | NW | NE | CE | SE | J-B |
| Itcha-Ilgachuz | – | <0.001 | < 0.001 | < 0.001 | < 0.001 | < 0.001 |
| NW | 0.086 (0.085-0.087) | – | < 0.001 | < 0.001 | < 0.001 | < 0.001 |
| NE | 0.101 (0.100-0.103) | 0.023 (0.023-0.024) | – | < 0.001 | < 0.001 | < 0.001 |
| CE | 0.132 (0.130-0.134) | 0.059 (0.058-0.060) | 0.062 (0.061-0.063) | – | < 0.001 | < 0.001 |
| SE | 0.096 (0.094-0.097) | 0.030 (0.029-0.030) | 0.028 (0.028-0.029) | 0.038 (0.037-0.038) | – | < 0.001 |
| J-B | 0.149 (0.147-0.151) | 0.072 (0.071-0.074) | 0.070 (0.069-0.071) | 0.051 (0.050-0.042) | 0.067 (0.066-0.068) | – |

**Table 3.** Observed (*H_o_*) heterozygosity, expected (*H_e_*) heterozygosity values and associated standard deviations, and *F_IS_* for inferred clusters at level 4 – 7 of our hierarchical analysis of woodland caribou in western Canada. Cluster labels follow the abbreviations in Table 1 and levels presented in Figure 4.

| Level/Cluster | $H_o$ | $H_e$ | $F_{IS}$ |
| --- | --- | --- | --- |
| **Level 1** |  |  |  |
| Itcha-Ilgachuz | $0.329 \pm 0.173$ | $0.327 \pm 0.166$ | <0.001 |
| All other subpopulations | $0.383 \pm 0.095$ | $0.409 \pm 0.099$ | 0.063 |
| **Level 2** |  |  |  |
| Itcha-Ilgachuz | $0.329 \pm 0.173$ | $0.327 \pm 0.166$ | <0.001 |
| Northern Cluster | $0.387 \pm 0.099$ | $0.407 \pm 0.101$ | 0.051 |
| Southern Cluster | $0.378 \pm 0.106$ | $0.399 \pm 0.108$ | 0.053 |
| **Level 3** |  |  |  |
| Itcha-Ilgachuz | $0.329 \pm 0.173$ | $0.327 \pm 0.166$ | <0.001 |
| NW | $0.380 \pm 0.112$ | $0.398 \pm 0.111$ | 0.048 |
| NE | $0.393 \pm 0.105$ | $0.405 \pm 0.103$ | 0.033 |
| CE | $0.389 \pm 0.110$ | $0.400 \pm 0.108$ | 0.030 |
| SE | $0.352 \pm 0.146$ | $0.369 \pm 0.140$ | 0.044 |
| J-B | $0.359 \pm 0.157$ | $0.360 \pm 0.142$ | 0.017 |

### Isolation-by-distance

Geographic distance explained 34.1% of the variation in Nei’s genetic distance across the sampled range (Mantel test, *p* < 0.001, Figure 7a). When testing for isolation-by-distance within each cluster at each hierarchical level up to level 3 (except the Itcha-Ilgachuz cluster which could not be tested as it only contained a single subpopulation), we identified significant or near-significant patterns of isolation-by-distance in all clusters (Figure 7).

**Figure 7.**
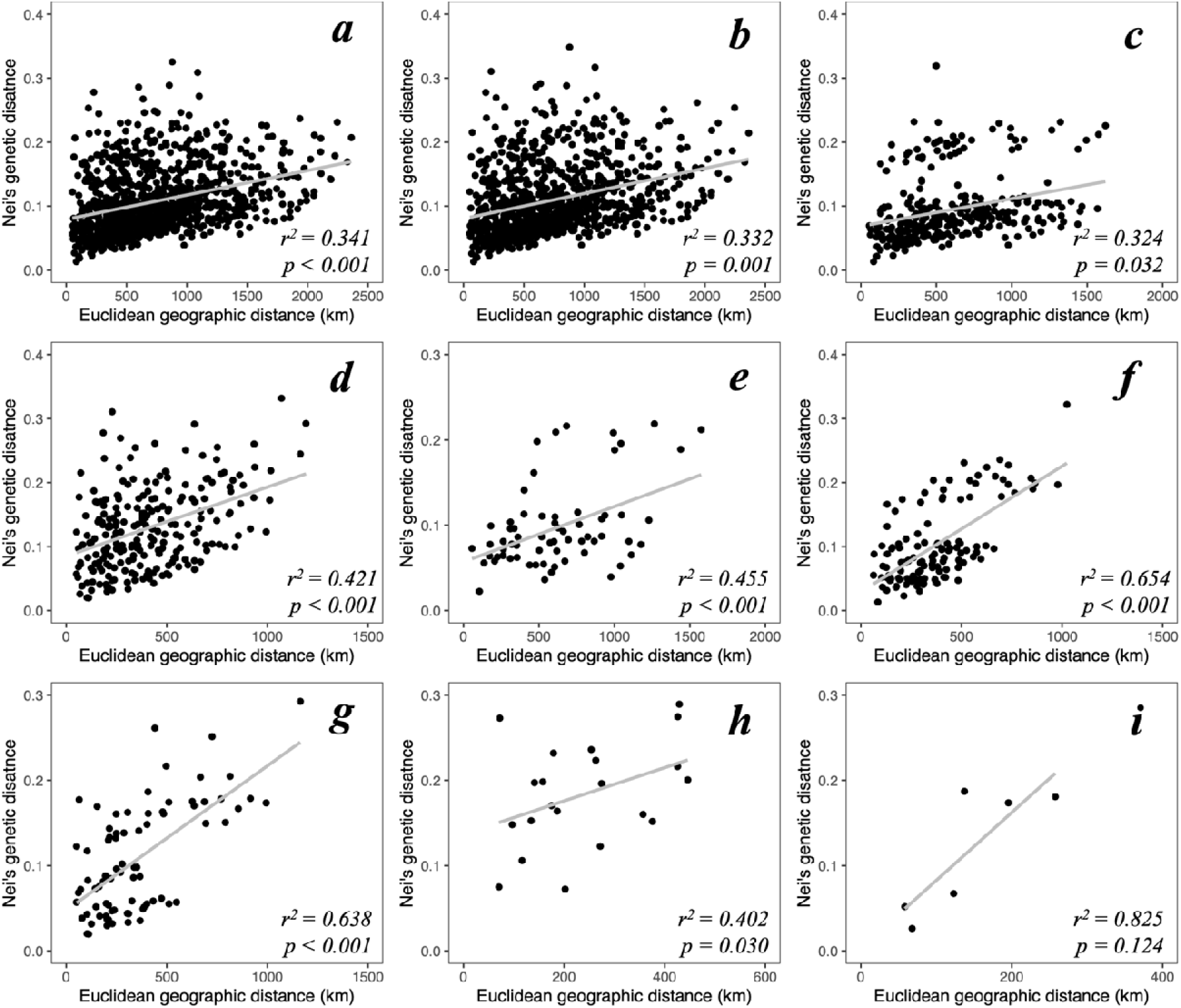
Nei’s genetic distances as a function of Euclidean geographic distance (km) between predefined subpopulations of woodland caribou in western Canada. Subpopulations were only included in the analysis if they had greater than five individuals genotyped. Panels depict analyses including (a) all subpopulations included in the study, or limited to those from (b) the main cluster at level 1, (c) the northern and (d) southern cluster at level 2, or the (e) northwestern, (f) northeastern, (g) central-eastern, (h) Jasper-Banff, and (i) southeastern clusters at level 3.

### Isolation by landscape features

When examining the geographic distribution of major genetic clusters, numerous landscape features appeared to be associated with the boundaries between genetic clusters. The Peace River and its drainage appeared to separate the northern and southern clusters at level 2; the northern extent of the Rocky Mountain Trench appeared to separate the northeastern and northwestern clusters; a combination of the North Thompson River and the headwaters of the Fraser River separate the Central-eastern and southeastern clusters; a combination of the Yellowhead Pass and Athabasca River separate the Central-eastern and Jasper-Banff clusters; and the Great Divide (the high elevation mountains along the border of Alberta and BC) appeared to separate the southeastern and Jasper-Banff clusters (Figures 1,4, and 5 [level 2]). Post hoc analyses indicated that four out of these five landscape features likely affected population structure. The Peace River and its drainage explained 5.9% of the variation in Nei’s genetic distance between subpopulations located north and south of the river in the main range (excluding Itcha-Ilgachuz; *p* < 0.001). North of the Peace River, separation across the Rocky Mountain Trench accounted for 16.3% of the variation in Nei’s genetic distance between the northeastern and northwestern subpopulations (*p* < 0.001). South of the Peace River, separation across the North Thompson River and headwaters of the Fraser River explained 41.9% of the variation in Nei’s genetic distance (*p* = 0.032), separation across the Yellowhead Pass and Athabasca River explained 43.0% of the variation in Nei’s genetic distance (*p* = 0.032), and South of the North Thompson River and Athabasca River, 41.5% of the variation in Nei’s genetic distance was attributed to separation east and west of the Great Divide; however, this result was only near-significant (*p* = 0.10).

## Discussion

In this study, we built on the preliminary work of Michalak (2023) to conduct a comprehensive hierarchical analysis of genetic diversity and variation in woodland caribou across western Canada. We found varying levels of genetic differentiation between subpopulations of woodland caribou in western Canada, explained by a combination of post-glacial recolonisation patterns, isolation-by-distance, and landscape features. Our clustering analysis identified hierarchical population structure ranging from *K=*2 at its highest level to *K=*31 at the lowest level, with *K=*6 appearing to be an appropriate broad scale grouping for capturing unique genetic diversity and informing conservation units. Patterns of genetic diversity across the landscape showed a tendency for higher diversity in the middle of the range, likely a result of secondary contact and historical hybridisation following post-glacial recolonisation. This study provides a comprehensive assessment of woodland caribou population structure in western Canada, which is essential to inform the delineation of conservation units for this species in the region.

### Broad-scale spatial genetic structure

Woodland caribou in western Canada were found to exhibit a multi-level hierarchical population genetic structure, consisting of two to six broad-scale genetic clusters. The greatest degree of separation was found between the Itcha-Ilgachuz subpopulation and all other subpopulations. We subsequently found an overarching north-south split among most subpopulations (all subpopulations when excluding Itcha-Ilgachuz) at the Peace River and its drainage which we attribute to a combination of glacial history in this region (Shafer et al. 2010) and separation by river system. Beyond this, six clusters broadly represent the Itcha-Ilgachuz subpopulation, northwestern BC, northeastern BC, central BC and Alberta, southeastern BC, and the Jasper-Banff region. This more localised population structure also appeared to result from limited gene flow across geographical features, particularly lowland habitats.

As previously mentioned, the most prominent segregation in the observed hierarchical population genetic structure is the separation of Itcha-Ilgachuz and all other subpopulations, which indicates Itcha-Ilgachuz possesses unique genetic variation which potentially results from either genetic drift or adaptation. In addition to the genetic differences reported here and elsewhere (Serrouya et al., 2012; Taylor et al., 2020), and this subpopulation exhibits differing behavioural tendencies (Hughes et al., 2025; Lamb et al., 2025) when compared to other subpopulations in western Canada. However, the underlying mechanisms responsible for the unique genetic signature remain unclear. Notably, Itcha-Ilgachuz caribou were distinct from both the northern and southern clusters at level 2, representative of the BEL and NAL ancestries. When comparing *F_ST_*distances between Itcha-Ilgachuz and other clusters, Itcha-Ilgachuz appears to be most similar to the northwestern cluster, likely explained by its geographic proximity to other subpopulations within this cluster. However, contrary to this, previous studies have identified the subpopulation to be most similar and share ancestry with subpopulations in central and southeastern BC (Taylor et al., 2021, 2024). One explanation for the separation of Itcha-Ilgachuz is that the subpopulation has been historically isolated (Taylor et al., 2020) and genetic drift has led to substantial genetic differentiation from other subpopulations. Alternatively, Itcha-Ilgachuz may have been recolonised by individuals from a cryptic glacial refugia, either located on the coast (Shafer et al., 2010, 2011) or in high altitude ice-free nunataks which were present in this region (Ryder et al., 2007). While further research will be needed to clarify this, our results suggest that Itcha-Ilgachuz is on its own evolutionary trajectory, likely with its own unique adaptive potential and hence should be managed and conserved as such.

In the main range of the subpopulations (all subpopulations other than Itcha-Ilgachuz) we identified a north-south split at the Peace River, representing the two glacial ancestries known to be present in this region. We observed a gradual rather than an abrupt shift from one ancestry to the other, and a low *F_ST_* distance between the two clusters, indicative of a hybrid zone between the two lineages. This finding aligns with studies in other species (Shafer et al., 2010) and other studies of woodland caribou in this region (Cavedon et al., 2022a; Cavedon et al., 2022b; Serrouya et al., 2012; Taylor et al., 2021; Yannic et al., 2014). Furthermore, this north-south divide is also reflected in some behavioural differences (Apps et al. 2001; Hughes et al., 2025; Lamb et al. 2025; Theoret et al. 2022). Interestingly, individuals from the boreal region, thought to stem from the NAL (Cavedon et al., 2022a; Cavedon et al., 2022b), grouped within the northern (BEL ancestry) cluster. This suggests that contemporary gene flow between boreal and more northern subpopulations to their west may have weakened historical differentiation between these groups.

Major genetic clusters appeared to be delineated by prominent features of the landscape. We found major lowland habitat features to delineate population structure between some clusters in our study range similar to other alpine adapted species (Deakin et al., 2020; Fedy et al., 2008; Sim et al., 2019). Two sub-clusters were identified within the main northern cluster; a northeastern and northwestern cluster. These clusters were defined by and likely are a result of reduced gene flow across the northern Rocky Mountain Trench. However, extremely low *F_ST_* values observed between these clusters suggest long-range movements of caribou in this region (Watters & DeMars, 2017) may lessen the effect of the barrier. We found the boundary between the central-eastern cluster and the southeastern cluster delineated by the North Thompson River and headwaters of the Fraser River west of The Great Divide; a boundary also observed in behavioural patterns (Hughes et al., 2025). East of the Great Divide the central-eastern cluster and Jasper-Banff clusters is delineated by the Yellowhead Pass and Athabasca River. Conversely, high elevation habitats along the Great Divide appeared to potentially separate the south-eastern and Jasper-Banff clusters, similar to other species where high alpine and mountainous habitats limit dispersal (Ghaedi et al., 2021; Machado et al., 2018; Rueness et al., 2003; Zalewski et al., 2009). However, this result was only near significant when tested with a partial Mantel test, likely due to our limited sample sizes in this analysis and region. It should be recognised that despite the apparent large effects of these features on genetic distance between major clusters, up to ∼43% in some cases, the total genetic distance between clusters at this level and among subpopulations in general tended to be low. These results further highlight how features of the landscape can influence gene flow and in turn population genetic structure, whether these features are energetically expensive habitats to traverse (Olah et al., 2017; Pérez-Espona et al., 2008), impassible or near impassible barriers (Epps et al., 2005; Hapeman et al., 2011; Rueness et al., 2003), or patches of undesirable habitat (Adams & Burg, 2015; Deakin et al., 2020).

### Fine-scale spatial structure

Spatial genetic structure was characterised by a pattern of isolation-by-distance, where geographic distance explained approximately a third of genetic distance between all pairs of subpopulations, likely resulting from both historical isolation and contemporary gene flow. The pattern of isolation-by-distance across the study area is lower than observed in other habitat specialised ungulates in this region, such as bighorn sheep (*Ovis canadensis canadensis*) (Deakin *et al*. 2020; Forbes and Hogg, 1999) and mountain goats (*Oreamnos americanus*) (Shafer et al., 2011), likely because woodland caribou are highly mobile and occupy highly fragmented habitats (Maltman et al., 2024), which may weaken the signal of isolation-by-distance.

Ultimately, our hierarchical population structure analysis identified many sub-clusters. These appeared to result from multiple causes including landscape features, geographic distance, and behaviour. For example, in the northeastern cluster we observed a split between Northern Mountain and Boreal individuals (SARA, 2012b, 2012a). Other breaks at lower hierarchical levels may be due to other landscape features, which are difficult to detect due to the numerous valleys, waterways, and habitats present in this highly heterogenous landscape. Overall, we found a total of 31 subpopulation level clusters, suggesting that not all the 45 pre-defined subpopulations are genetically distinct. This indicates that in some cases multiple predefined subpopulations may function as one or have become geographically isolated relatively recently.

### Patterns of genetic diversity

We observed patterns of genetic diversity concordant with the history of post-glacial recolonisation of the area (Hewitt, 2004; Shafer et al., 2010, 2011). Typically, it is expected that genetic diversity should decrease with distance from source populations due to the founder effect (Frankham, 1997), but here we found genetic diversity to be lower in the northwest and southeast and highest in the middle of the sampled range. The elevated genetic diversity in the middle of our sampling range is likely due to hybridisation following secondary contact between the two historical lineages in this region (Barton & Hewitt, 1985; Canestrelli et al., 2010; Cavedon et al., 2022a; McDevitt et al., 2009). Although subpopulations in the central part of this range are among some of the most at risk (Lamb et al., 2024), their elevated genetic diversity means they may also possess more adaptive potential and thus be more resilient to future environmental and habitat changes.

### Conservation implications

As suggested by the preliminary analyses of Michalk (2023), boundaries between major genetic clusters do not fully align with currently recognised conservation units in the study region. While four of the six major genetic clusters identified somewhat resemble existing DU and SARA classification schemes (COSEWIC, 2011; SARA, 2012b, 2012a, 2014), boundary shifts and additional units are required to more appropriately delineate unique genetic variation in the region (Figure 1). The boundary between the Boreal and the Northern Mountain DU and SARA units should be shifted westward to the Northern Rocky Mountain Trench to include northern subpopulations east of this lowland system, forming northeastern and northwestern clusters. In the southern DU and SARA units, boundary redefinition should also be considered to reflect the differences between the central-eastern, southeastern, and newly identified Jasper-Banff clusters; with a north-south split delineated by the North Thompson River, Fraser River headwaters, and Athabasca River, and an east-west split south of this along the Great Divide (separating out the Jasper-Banff region). Additionally, our analyses revealed that the Itcha-Ilgachuz subpopulation forms a unique genetic cluster, highly distinct from all others, and may warrant being managed as such (Figure. 1).

All broad-scale genetic clusters (levels 1 – 3) exhibit significant genetic differentiation, supporting their classification as distinct conservation units (COSEWIC, 2011; Fraser & Bernatchez, 2001; Hoelzel, 2023; Muir et al., 2021). However, as highlighted by Hoelzel (2023), estimates of genetic differentiation with vast numbers of markers may overstate the significance of these differences. While gene flow between major clusters appears low, genetic differences should be considered in the context of overall variation, especially when factoring in behavioural distinctions (COSEWIC, 2011; SARA, 2012b, 2012a, 2014; Theoret et al., 2022), and known differences in glacial ancestry (McDevitt et al., 2009; Taylor et al., 2021; Yannic et al., 2014). Hence, we propose that the six major genetic clusters identified in our study represent six conservation units, which contain both unique and irreplaceable genetic variation. Thus, highlighting the need for tailored conservation measures to preserve these evolutionary legacies.

## Conclusion

In this study, we built on the preliminary work of Michalak (2023) to characterised genetic diversity across woodland caribou range in western Canada. We identified a hierarchical population structure composed of multiple populations and subpopulations, which is best described by six genetic clusters: the northeastern, northwestern, central-eastern, southeastern, the Jasper-Banff, and Itcha-Ilgachuz clusters. This structure is indicative of post-glacial recolonisation, particularly a north-south split, with patterns of hybridisation shaping genetic diversity across the landscape. We suggest that current conservation units for woodland caribou in this region should be revised to account for this information. This study exemplifies how wide-ranging, mobile species can exhibit intricate population genetic structure, especially those with complex natural histories occupying highly heterogeneous landscapes. In such cases, range-wide examination of genetic data is crucial for identifying or refining conservation units.

## Acknowledgements

We extend our sincere gratitude to the government employees, students, First Nations, the Nîkanêse Wah tzee Stewardship Society, and other community members whose efforts in sample and data collection were instrumental in making this work possible. This research was financially supported by the Government of British Columbia, the Canadian Wildlife Service, Parks Canada, and Natural Sciences and Engineering Research Council of Canada Grants to JP and MM. AM was supported by an Alberta Graduate Excellence Scholarship.

## Data Accessibility

This study is based on data provided or generated under research agreements between the University of Calgary and government partners which forbid public release of the raw data. Access to data may be granted following completion of data sharing agreements with the appropriate government agencies.

## Author Contributions

**Samuel Deakin**: Writing – original draft, Writing – final draft, review & editing, Visualization, Methodology, Investigation, Formal analysis, Conceptualization, Data curation. **Anita Michalak:** Writing – original draft, review & editing, Visualization, Methodology, Investigation, Formal analysis, Conceptualization, Data curation. **Maria Cavedon**: Writing – review & editing, Methodology, Formal analysis, Conceptualization. **Charlotte Bourbon**: Writing – review & editing, Methodology. **Margaret Hughes**: Writing – review & editing, Methodology. **Lalenia Neufeld**: Writing – review & editing, Conceptualization, Methodology, Funding acquisition, Project administration, Data curation. **Agnès Pelletier**: Writing – review & editing, Methodology, Conceptualization. **Jean Polfus**: Writing – review & editing, Conceptualization, Funding acquisition, Project administration. **Helen Schwantje**: Writing – review & editing, Methodology, Conceptualization, Funding acquisition, Project administration, Data curation. **Robin Steenweg:** Writing – review & editing, Funding acquisition, Conceptualization, Project administration. **Caeley Thacker**: Writing – review & editing, Methodology, Conceptualization, Funding acquisition, Project administration, Data curation. **Madeline Trottier**: Writing – review & editing, Data curation. **Marco Musiani**: Writing – review & editing, Supervision, Funding acquisition, Conceptualization, Methodology, Project administration. **Jocelyn Poissant**: Writing – review & editing, Supervision, Funding acquisition, Conceptualization, Methodology, Project administration.

## Notes

### Competing Interest Statement

The authors have declared no competing interest.

