## Supplemental Information for "Glacial history and landscape features shape the hierarchical population genetic structure of woodland caribou (*Rangifer tarandus caribou*) in western Canada"

**<u>Supplementary Materials</u>**

**Supplementary Information 1** - Script used to align reads to new chromosomal level genome

```
bowtie2-build -f -q GCA_019903745.2_ULRtarCaribou_2v2_genomic.fasta
GCA_019903745.2_ULRtarCaribou_2v2_genomic.bwt2ref

bowtie2 --phred33 --sensitive -N 1 -I 0 -X 500 -t -p 16 -x
GCA_019903745.2_ULRtarCaribou_2v2_genomic.bwt2ref -f
RangiferSNP63k_probe_seqs_TOPGENOMICSEQ.fasta -S
RangiferSNP63k_probe_seqs_TOPGENOMICSEQ.sam

samtools faidx GCA_019903745.2_ULRtarCaribou_2v2_genomic.fasta
samtools view -bt GCA_019903745.2_ULRtarCaribou_2v2_genomic.fasta.fai
RangiferSNP63k_probe_seqs_TOPGENOMICSEQ.sam >
RangiferSNP63k_probe_seqs_TOPGENOMICSEQ.bam
samtools view -b -F4 RangiferSNP63k_probe_seqs_TOPGENOMICSEQ.bam >
RangiferSNP63k_probe_seqs_TOPGENOMICSEQ.mapped.bam
samtools sort RangiferSNP63k_probe_seqs_TOPGENOMICSEQ.mapped.bam >

RangiferSNP63k_probe_seqs_TOPGENOMICSEQ.mapped.sorted.bam
```

**Table S1.** Models fitted to investigate the association of census size and latitude with expected heterozygosity. *df* = degrees freedom, $AIC_C$ = Akaike Information Criterion corrected for small sample size, $\Delta AIC_C$ = delta $AIC_C$.

| Model | Fixed effects | Log-likelihood | df | $r^2$ | $AIC_C$ | $\Delta AIC_C$ | $AIC_C$ weight |
|---|---|---|---|---|---|---|---|
| Model 11 | latitude + latitude$^2$ + log census size | 93.89 | 5 | 0.56 | -176.45 | 0 | 0.669 |
| Model 10 | latitude + latitude$^2$ | 91.68 | 4 | 0.5 | -174.59 | 1.86 | 0.199 |
| Mode l9 | latitude + latitude$^2$ + census size | 92.27 | 5 | 0.52 | -173.16 | 3.29 | 0.129 |
| Model 8 | latitude + log census size | 86.21 | 4 | 0.32 | -163.64 | 12.81 | 0.001 |
| Model 7 | latitude$^2$ + log census size | 86 | 4 | 0.31 | -163.24 | 13.21 | 0.001 |
| Model 6 | latitude | 83.9 | 3 | 0.23 | -161.43 | 15.02 | <0.001 |
| Model 5 | latitude$^2$ | 83.66 | 3 | 0.22 | -160.94 | 15.51 | <0.001 |
| Model 4 | log census size | 83.14 | 3 | 0.19 | -159.91 | 16.54 | <0.001 |
| Model 3 | latitude + census size | 83.96 | 4 | 0.23 | -159.15 | 17.3 | <0.001 |
| Model 2 | latitude$^2$ + census size | 83.72 | 4 | 0.22 | -158.67 | 17.78 | <0.001 |
| Model 1 | census size | 80.1 | 3 | 0.04 | -153.83 | 22.62 | <0.001 |

**Table S2.** Parameter estimates for the best fitting model describing expected heterozygosity in woodland caribou subpopulations in western Canada.

| Parameter | Coefficient | SE | *t* | *p* |
|---|---|---|---|---|
| Intercept | -5.440 | 1.350 | -4.024 | <0.001 |
| Latitude | 0.200 | 0.049 | 4.200 | <0.001 |
| Latitude^2 | -0.002 | <0.001 | -4.130 | <0.001 |
| Log census size | 0.005 | 0.003 | 2.043 | 0.054 |

**Table S3.** 31final subpopulation level clusters of woodland caribou subpopulations in western Canada. Numbers in each cell denotes the number of individuals from that putative subpopulation in each cluster.

| Region | II | NW | | | | | | | | NE_W | | | NE_E | | | | CE_N | | CE_S | | | | | | J-B | | SM_S | | | | |
|---|---|---|---|---|---|---|---|---|---|---|---|---|---|---|---|---|---|---|---|---|---|---|---|---|---|---|---|---|---|---|---|
| Cluster | 1 | 2 | 3 | 4 | 5 | 6 | 7 | 8 | 9 | 10 | 11 | 12 | 13 | 14 | 15 | 16 | 17 | 18 | 19 | 20 | 21 | 22 | 23 | 24 | 25 | 26 | 27 | 28 | 29 | 30 | 31 |
| II | 75 | | | | | | | | | | | | | | | | | | | | | | | | | | | | | | |
| AT | | | | | # | | | | | | | | | | | | | | | | | | | | | | | | | | |
| CA | | | | | | 5 | | | | | | | | | | | | | | | | | | | | | | | | | |
| HO | | | | | | | | 4 | | | | | | | | | | | | | | | | | | | | | | | |
| LK | | | | | | | 3 | | | | | | | | | | | | | | | | | | | | | | | | |
| LR | | | | | | | 6 | | | | | 1 | | | | | | | | | | | | | | | | | | | |
| TS | | | | | | | | 7 | | | | | | | | | | | | | | | | | | | | | | | |
| CH | | | # | 6 | | | | 1 | | | | 3 | | | | | | | | | | | | | | | | | | | |
| TK | | 28 | 1 | | | | | | 4 | | | | | | | | | | | | | | | | | | | | | | |
| TW | | | | | | | | | | | | | | | | | | | | | | | | | | | | | | | |
| WO | | | 2 | 34 | | | | 1 | | | | | | | | | | 1 | | | | | | | | | | | | | |
| GH | | | | | | | | | | | 22 | | | | | | | | | | | | | | | | | | | | |
| CL | | | | | | | | | | | | | | | 8 | 1 | | | | | | | | | | | | | | | |
| CC | | | | | | | | | | | | | 20 | | | 1 | | | | | | | | | | | | | | | |
| HA | | | | | | | | | | | | | | | | 1 | | | | | | | | | | | | | | | |
| MX | | | | | | | | | | 1 | | | | | | 7 | | | | | | | | | | | | | | | |
| SS | | | | | | | | | | | | | | 1 | | 21 | | | | | | | | | | | | | | | |
| WFN | | | | | | | | | | 1 | 1 | | | 4 | | 1 | | | | | | | | | | | | | | | |
| EW | | | | | | | | | | | 3 | | | | | | | | | | | | | | | | | | | | |
| FI | | | | | | | | | | | 5 | | | | | | | | | | | | | | | | | | | | |
| FR | | | | | | | | 3 | | | | 3 | | | | | | | | | | | | | | | | | | | |
| GA | | | | | | | | | | | | 5 | | | | | | | | | | | | | | | | | | | |
| MU | | | | | | | | | | 26 | | 2 | | | | | | | | | | | | | | | | | | | |
| PM | | | | | | | | | | | 39 | | | | | | | | | | | | | | | | | | | | |

| | | | | | | | | | | | | | | | | | | | | | | | | | | | | | | | |
|---|---|---|---|---|---|---|---|---|---|---|---|---|---|---|---|---|---|---|---|---|---|---|---|---|---|---|---|---|---|---|---|
| ALP | | | | | | | | | | | | | | | | | | | | | 2 | | 12 | | | | | | | | |
| KS | | | | | | | | | | 2 | 3 | | | | | 10 | 1 | | | | | | | | | | | | | | |
| KZ | | | | | | | | | | | | | | | | 20 | | | | | | | | | | | | | | | |
| NA | | | | | | | | | | | | | | | | 1 | 2 | | | | 1 | | | | | | | | | | |
| QI | | | | | | | | | | | | | | | | | 15 | | | | 1 | | | | | | | | | | |
| RPC | | | | | | | | | | | | | | | | | | | | 9 | 1 | | | | | | | | | | |
| BR | | | | | | | | | | | | | | | | | | 4 | | | | | | | | | | | | | |
| GR | | | | | | | | | | | | | | | | | | | | | | | | | | | | | | | 3 |
| HR | | | | 1 | | | | | | | | | | | | | 24 | | | | 34 | | | | | | | | | | |
| NL | | | | | | | | | | | | | | | | | | | | | 2 | | | | | | | | | | |
| NC | | | | | | | | | | | | | | | | | | | | | 2 | 21 | | | | | | | | | |
| WGS | | | | | | | | | | | | | | | | | | 3 | 15 | | | | | | | | | | | | |
| CK | | | | | | | | | | | | | | | | | | | | | | | | | | | 6 | | | | |
| CN | | | | | | | | | | | | | | | | | | | | | | | | | | 31 | | | | | 1 |
| CS | | | | | | | | | | | | | | | | | | | | | | | | | | | | | | 1 | |
| PS | | | | | | | | | | | | | | | | | | | | | | | | | | | | | 2 | 2 | |
| SK | | | | | | | | | | | | | | | | | | | | | | 1 | | | | | | | | 1 | |
| BNP | | | | | | | | | | | | | | | | | | | | | | | | | 1 | | | | | | |
| BZ | | | | | | | | | | | | | | | | | | | | | | | | | 5 | | | | | | |
| ML | | | | | | | | | | | | | | | | | | | | | | | | 4 | 5 | | | | | | |
| TQ | | | | | | | | | | | | | | | | | | | | | | | | 20 | 1 | | | | | 1 | |
| Total | 75 | 28 | # | 41 | # | 5 | 9 | 16 | 4 | 28 | 72 | 17 | 20 | 5 | 8 | 32 | 31 | 43 | 7 | 15 | 9 | 43 | 22 | 12 | 24 | 12 | 31 | 6 | 2 | 5 | 4 |

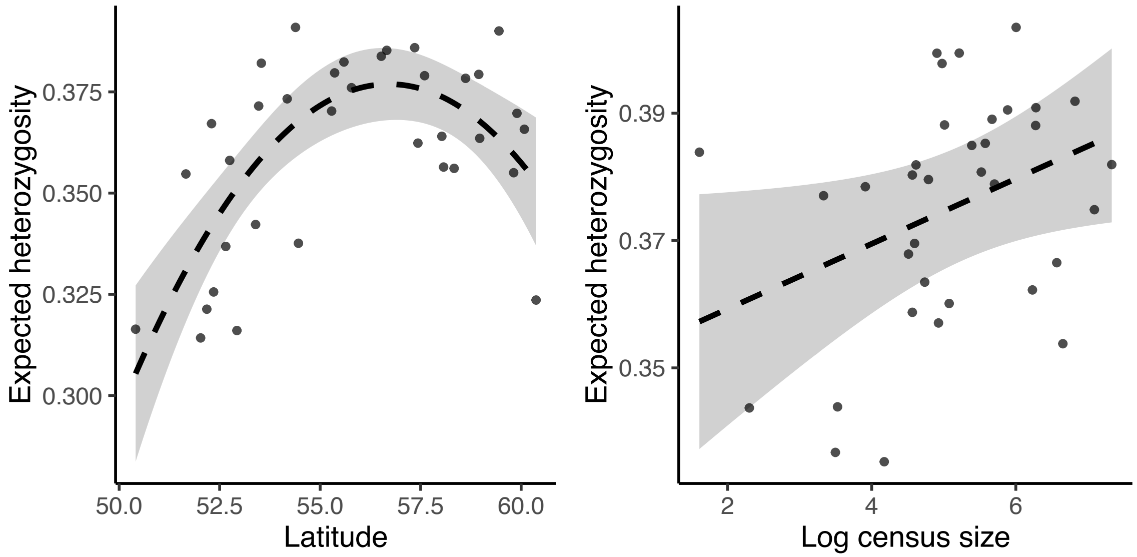


**Figure S1.** Predicted associations between a) latitude (degrees North) and b) log-transformed census size and expected heterozygosity of woodland caribou subpopulations in western Canada from a linear mixed model. Plots generated using the R *visreg* function. Points show changes in response while holding all other variables constant. Grey area depicts 95% confidence intervals of predicted associations.

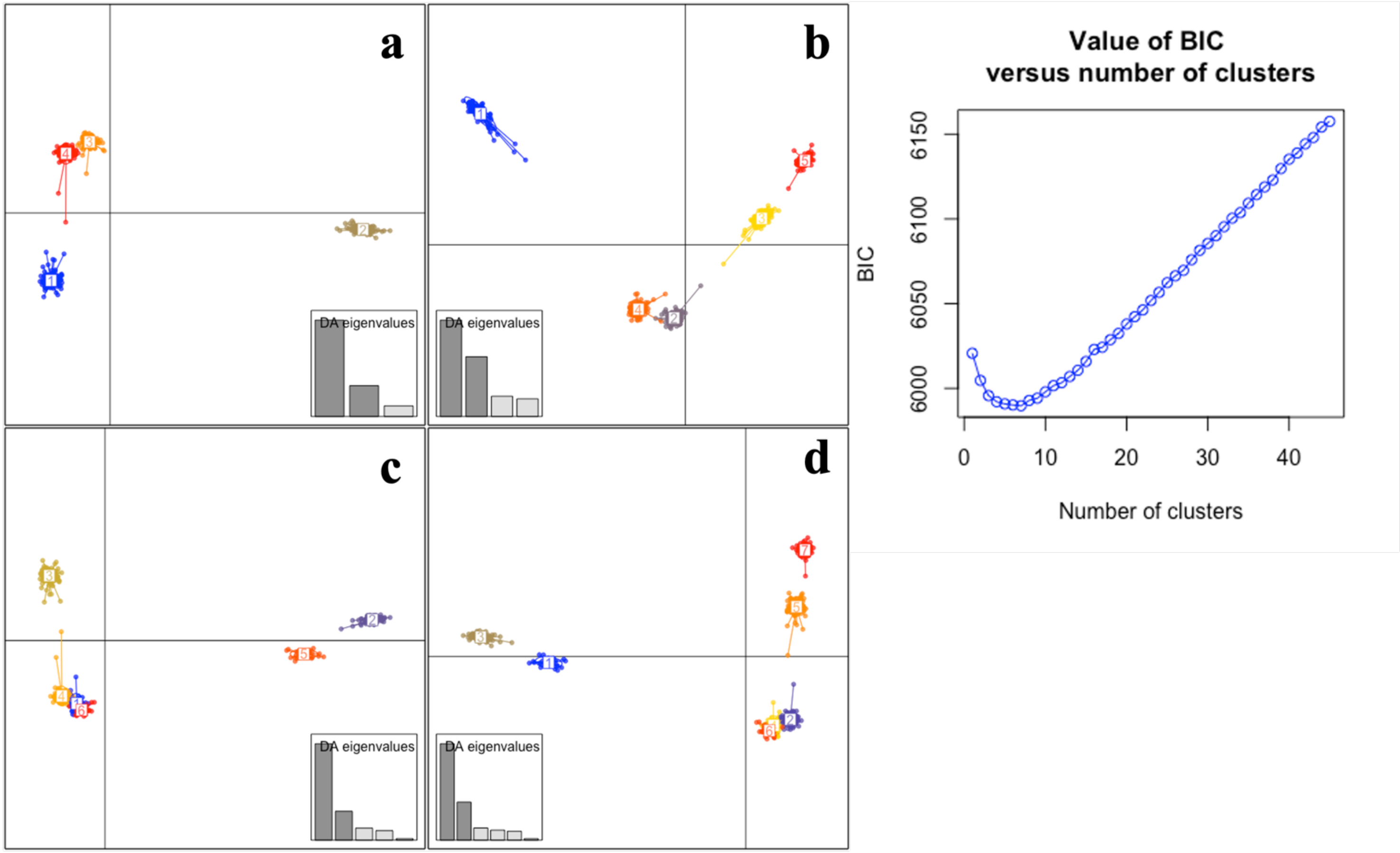


**Figure S2.** Scatterplot of Discriminant Analysis of Principal Components (DAPC) results retaining from 44 principal components and indicating a separation of individuals into four (a), five (b), six (c), and seven (d) clusters.

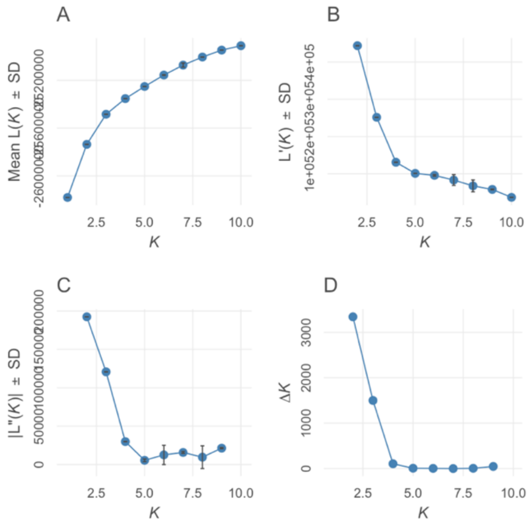


**Figure S3**. Log likelihood plots generated by pophelper for level 1.

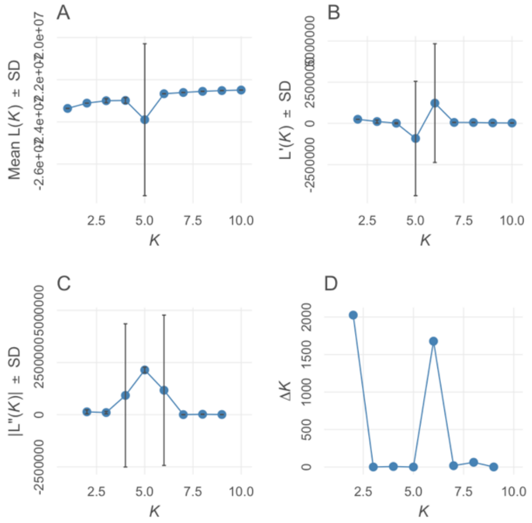


**Figure S4**. Log likelihood plots generated by pophelper for level 2.

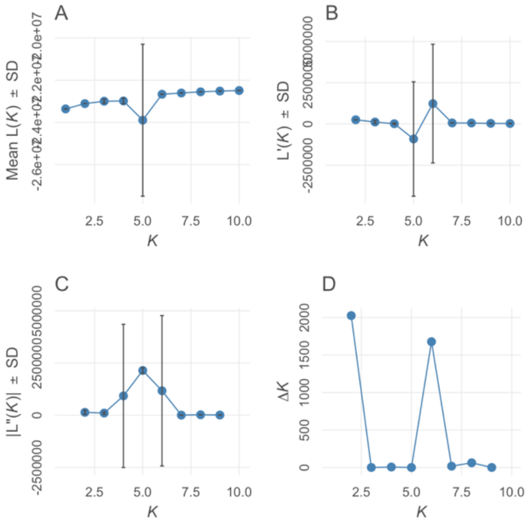


**Figure S5**. Log likelihood plots generated by pophelper for northern cluster level 3

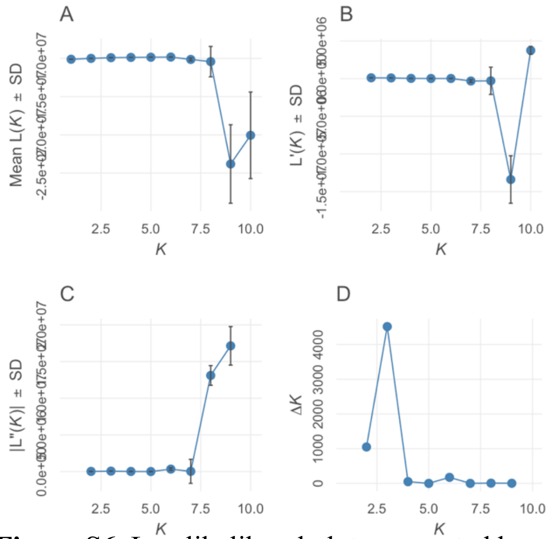


**Figure S6**. Log likelihood plots generated by pophelper for southern cluster level 3.

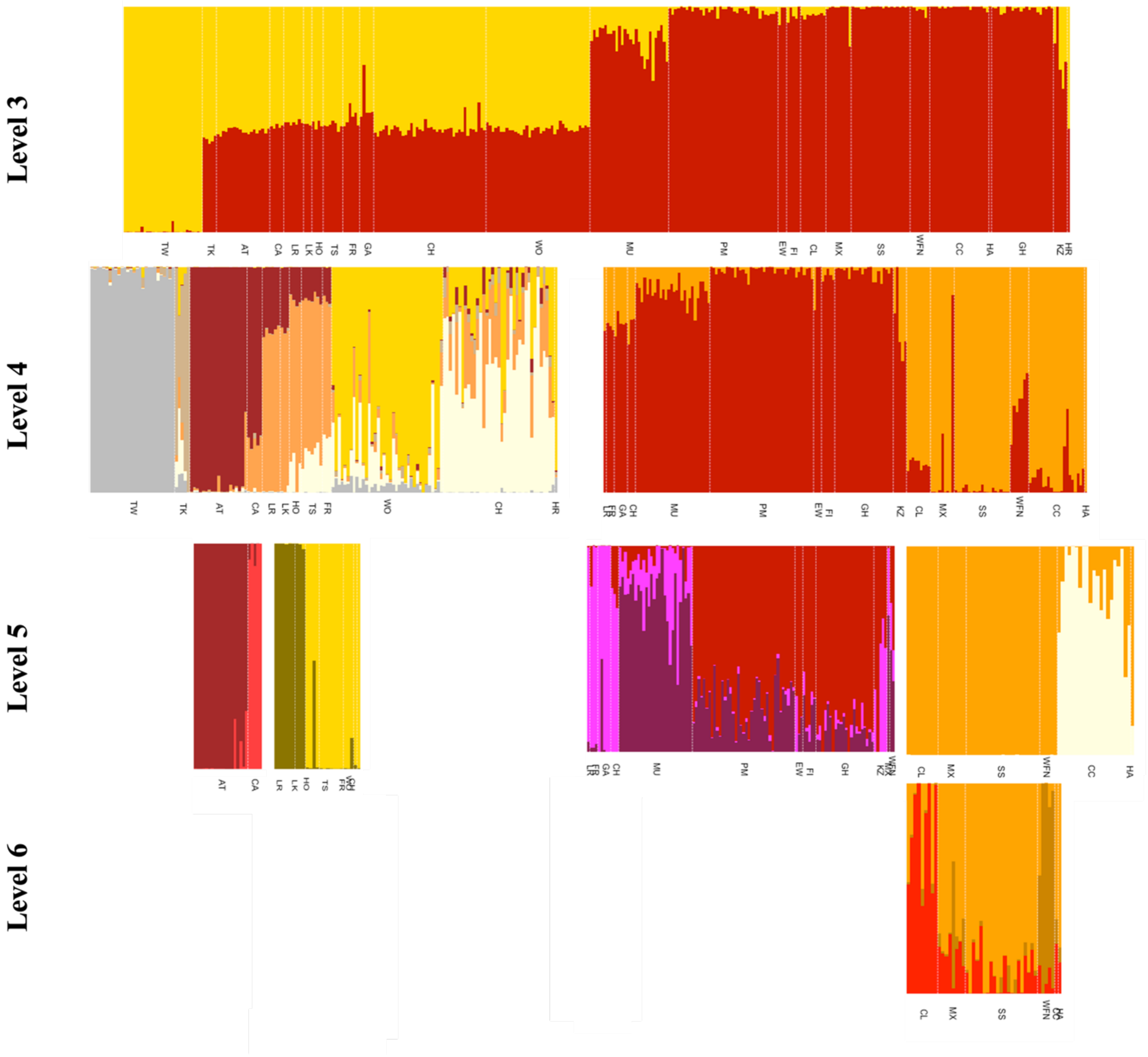


**Figure S7.** Admixture plots for woodland caribou (*Rangifer tarandus caribou*) included in subpopulation structure analysis from the northwestern cluster (*n*=337). Labels correspond to individuals' putative subpopulations. Dashed boxes indicate what we consider to be resolved subpopulations.

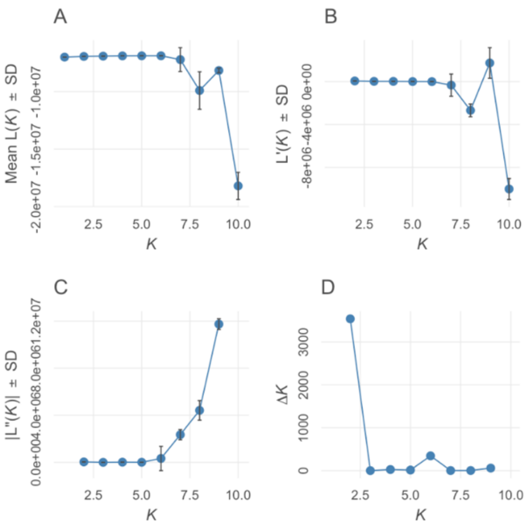


**Figure S8**. Log likelihood plots generated by pophelper for northeastern cluster level 4

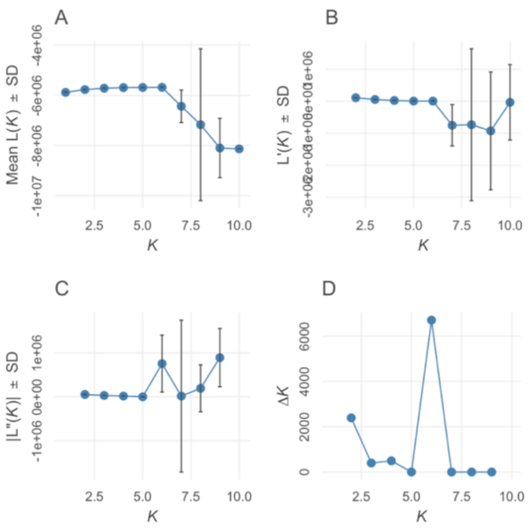


**Figure S9**. Log likelihood plots generated by pophelper for northwestern cluster level 4

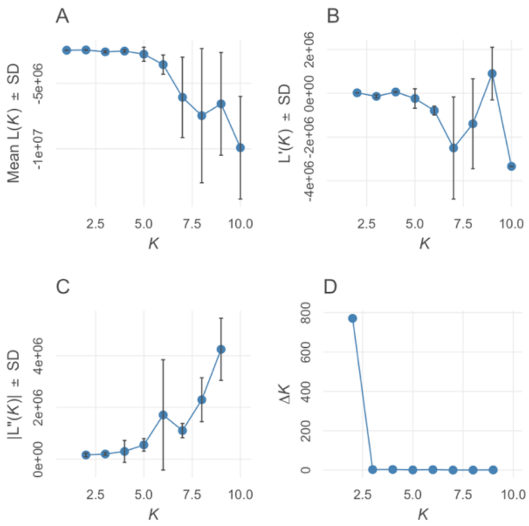


**Figure S10**. Log likelihood plots generated by pophelper for Boreal cluster level 5.

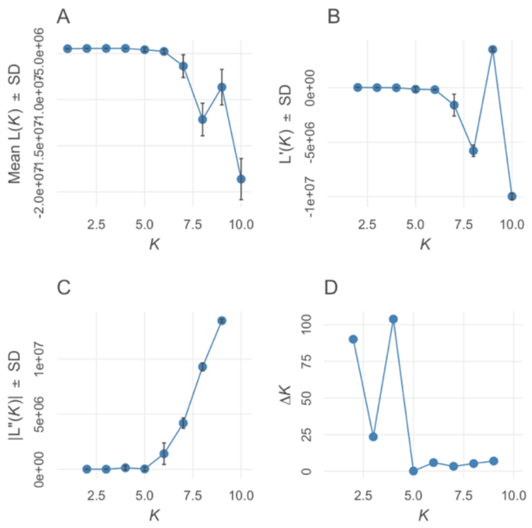


**Figure S11**. Log likelihood plots generated by pophelper for northeastern slopes cluster level 5.

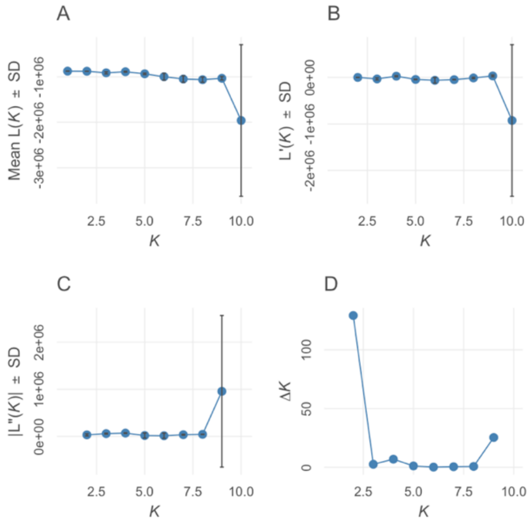


**Figure S12**. Log likelihood plots generated by pophelper for Atlin and Carcross cluster level 5.

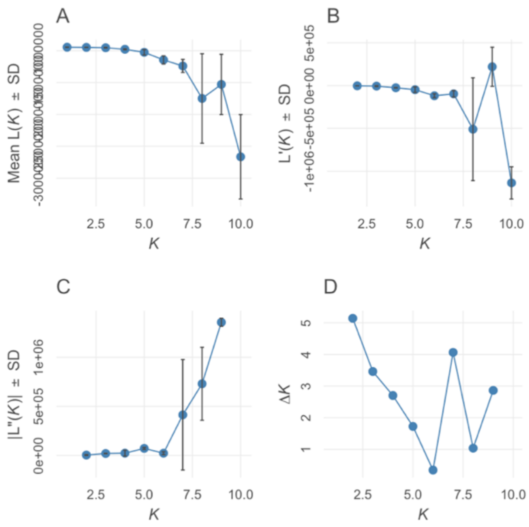


**Figure S13**. Log likelihood plots generated by pophelper for Horseranch, Tsenaglode, Level-Kawdy, Little Rancheria, Frog, Wolverine, and Chase cluster level 5.

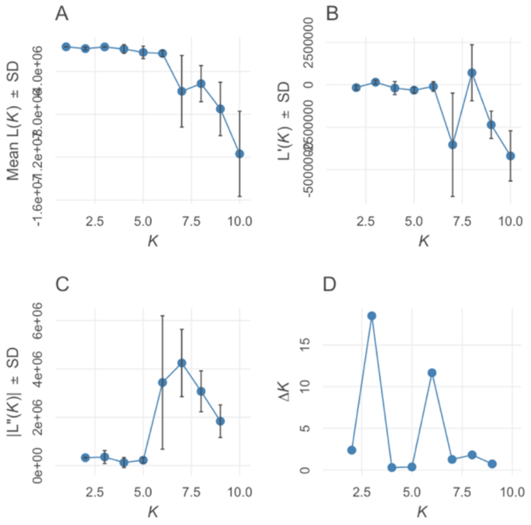


**Figure S14**. Log likelihood plots generated by pophelper for the northern Boreal cluster level 6.

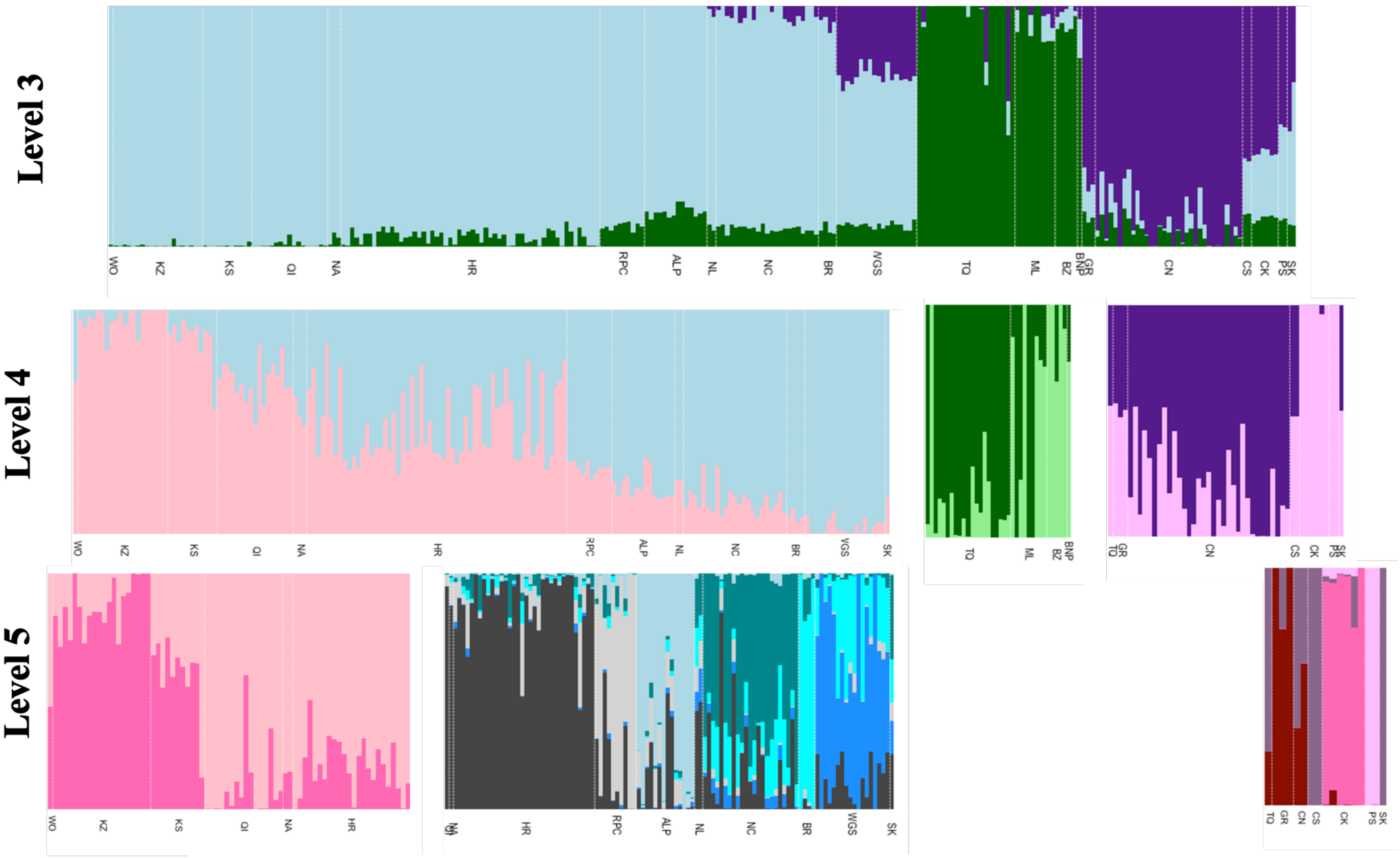


**Figure S15.** Admixture plots for woodland caribou (*Rangifer tarandus caribou*) included in subpopulation structure analysis from the central-eastern cluster (*n*=266). Labels correspond to individuals' putative subpopulations.

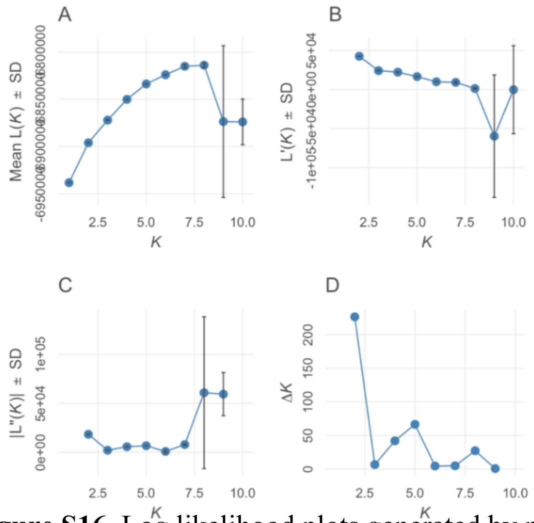


**Figure S16**. Log likelihood plots generated by pophelper for central-eastern cluster at level 4.

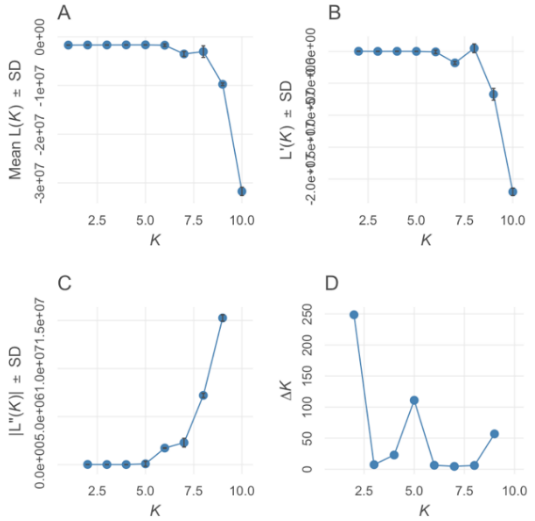


**Figure S17**. Log likelihood plots generated by pophelper for southeastern cluster at level 4.

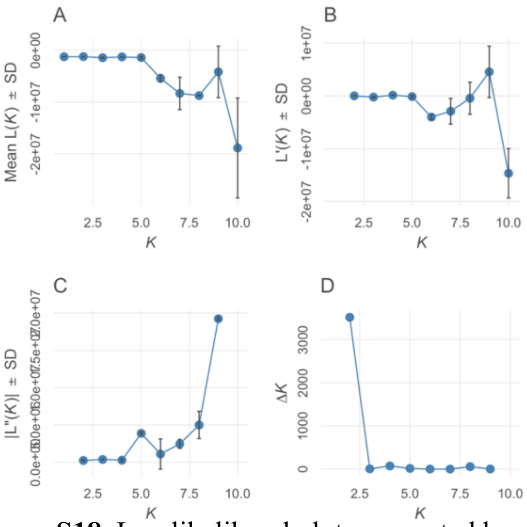


**Figure S18**. Log likelihood plots generated by pophelper for northern Jasper-Banff cluster at level 4.

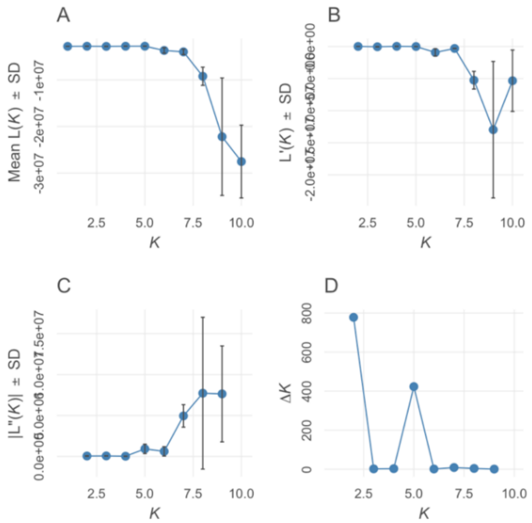


**Figure S19**. Log likelihood plots generated by pophelper for northern central-eastern cluster at level 4.

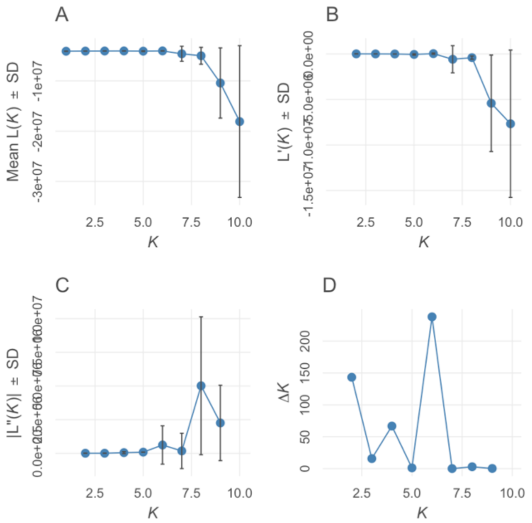

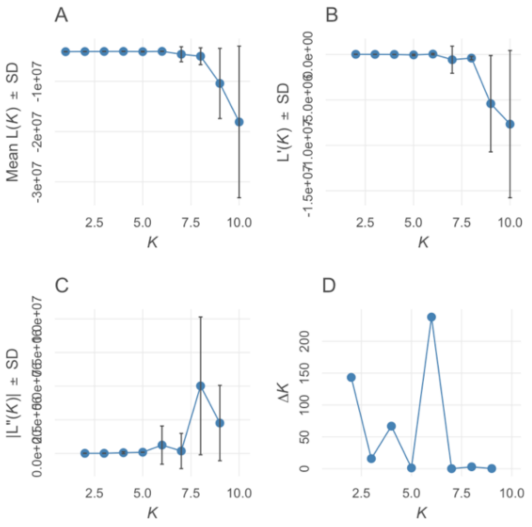


**Figure S20**. Log likelihood plots generated by pophelper for the southern central-eastern cluster at level 5.

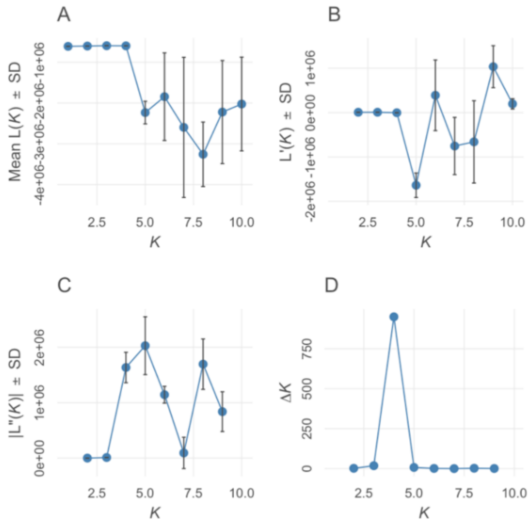


**Figure S21**. Log likelihood plots generated by pophelper for the southeastern cluster at level 5.

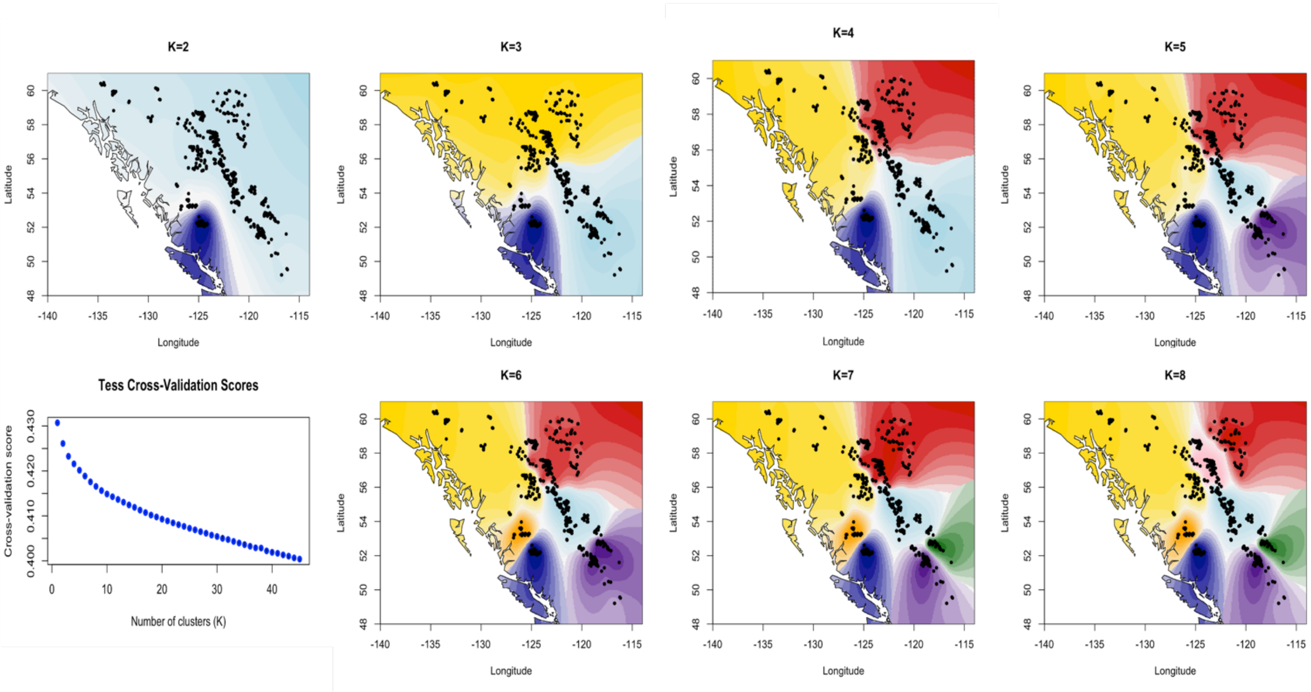


**Figure S22**. *TESS* cross validation scores and predictions of genetic and geographic clusters inferred from *TESS* $K = 2–8$ for 678 woodland caribou distributed across 45 pre-defined subpopulations in western Canada. Maps created using the *tess3r* package in R.

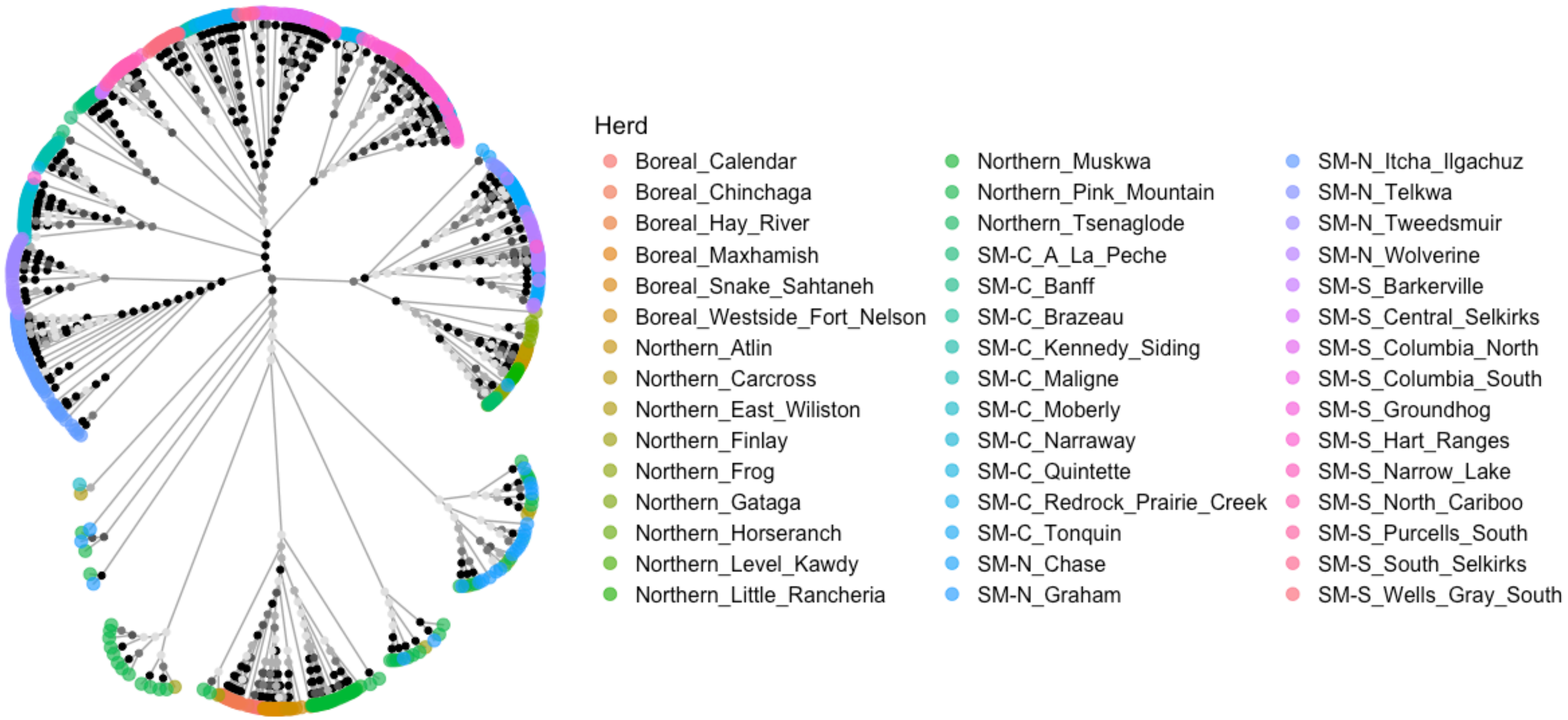


**Figure S23.** Neighbor-joining tree of woodland caribou sampled throughout British Columbia and part of Alberta. Branches represent individuals, with tip colours representing each individual's source subpopulation and SARA listing (SARA, 2014). Bootstrap values were estimated based on 1000 replicates and are represented on internal nodes as circles in five classes/shades of grey (in 20% increments, with the darkest circle representing the 81–100% class). Abbreviations: SM-N = Southern Mountain-Northern Group, SM-C = Southern Mountain-Central Group, and SM-S = Southern Mountain-Southern Group.
